# Context-independent scaling of neural responses to task difficulty in the multiple-demand network

**DOI:** 10.1101/2022.08.12.503813

**Authors:** Tanya Wen, Tobias Egner

## Abstract

The multiple-demand (MD) network is sensitive to many aspects of cognitive demand, showing increased activation with more difficult tasks. However, it is currently unknown whether the MD network is modulated by the context in which task difficulty is experienced. Using fMRI, we examined MD network responses to low, medium, and high difficulty arithmetic problems within two cued contexts, an easy versus a hard set. The results showed that MD activity varied reliably with the absolute difficulty of a problem, independent of the context in which the problem was presented. Similarly, MD activity during task execution was independent of the difficulty of the previous trial. Representational similarity analysis further supported that representational distances in the MD network were consistent with a context-independent code. Finally, we identified several regions outside the MD network that showed context-dependent coding, including the inferior parietal lobule, paracentral lobule, posterior insula, and large areas of the visual cortex. In sum, cognitive effort is processed by the MD network in a context-independent manner. We suggest that this absolute coding of cognitive demand in the MD network reflects the limited range of task difficulty that can be supported by the cognitive apparatus.

---

The multiple-demand (MD) network is a set of frontal and parietal brain regions whose responses scale with cognitive demands, exhibiting enhanced activity with increasing cognitive load or difficulty across a diverse set of tasks (Duncan and Owen 2000; Duncan 2010; Fedorenko et al. 2013; Duncan et al. 2020). To account for this broad association with cognitive demand, the MD network has been suggested to implement top down control to focus on the operations required for a current task, regardless of the precise nature of those operations (Erez and Duncan 2015; Jackson et al. 2017; Wen, Duncan, et al. 2020). However, as most previous studies were limited to a single experimental context in which difficulty was manipulated, a fundamental question about the relationship between MD network activity and cognitive effort remains unanswered: is MD network activity shaped by the context in which a given level of task difficulty is experienced?

On the one hand, several lines of research speak in favor of the possibility that difficulty or effort could be coded in a content-sensitive manner in the MD network. First, the MD network has been implicated in flexible coding of context-dependent task rules, producing distinct responses to the same input data, such as judging the dominant color or motion direction of a cloud of moving dots (Roy et al. 2010; Mante et al. 2013; Flesch et al. 2022). Thus, it seems plausible that the same level of task difficulty could produce different responses under different contexts in this network. Second, context-dependent coding – sometimes referred to as “range adaptation” – is commonly observed in value-based decisions and perceptual processing (Nieuwenhuis et al. 2005; Elliott et al. 2008; Carandini and Heeger 2011; Cheadle et al. 2014; Cox and Kable 2014; Palminteri et al. 2015; Murai et al. 2016; Bavard et al. 2018, 2021; Hunter and Daw 2021). For example, Nieuwenhuis et al. (2005) created two contexts in which participants would either always win or always lose money. Within each context, there were also three possible outcomes, worst (+0¢/-40¢), intermediate (+30¢/-20¢), and best (+60¢/-0¢). The authors found that activity in reward-sensitive areas scaled positively with outcome value (best > intermediate > worst) in each context, but that activity levels for the three outcomes were comparable between contexts, despite the large difference in the objective value of these outcomes. In other words, neural reward coding appears to be relative, such that an equivalent absolute value will elicit a greater response if it is a relatively good than if it is a relatively bad outcome in the current context.

Range adaptation in reward coding is relevant to the MD network because of its association with cognitive effort, whose recruitment is comonly conceptualized as directly proportional to prospective reward (e.g., Shenhav et al., 2013). Specifically, previous work has shown that MD regions may be divided into two closely coupled networks centered around the frontoparietal and cingular-opercular network (Dosenbach et al. 2008; Crittenden et al. 2016; Shashidhara et al. 2019), with the former consisting several distinct regions of the middle and posterior frontal cortex (mPFC and pMFC), posterior-dorsal lateral frontal cortex (pdLFC), and the intraparietal suclus (IPS), while the later consists of the anterior medial prefontal cortext (aMFC), anterior insula (AI), and dorsal anterior cingulate cortex (ACC). Recent studies have identified overlapping regions involved in cognitive effort and the anticipation and processing of reward in the MD network, especially in cingular-opercular regions (Prévost et al. 2010; Shashidhara et al. 2019), including the AI (Lallement et al. 2014; Chong et al. 2017) and ACC (Croxson et al. 2009; Vassena et al. 2014; Chong et al. 2017). In particular, the ACC has been proposed to play a critical role in allocating cognitive control based on the evaluation of the reward and costs that can be expected from a control-demanding task, referred to as the expected value of control (Shenhav et al. 2013). Furthermore, in effort discounting tasks, researchers have identified that frontoparietal regions track shared variance between subjective value and difficulty (Massar et al. 2015; Chong et al. 2017; Westbrook et al. 2019). In line with studies suggesting a close relationship between effort and reward of cognitive actions (Kool et al. 2010; Otto and Vassena 2021), it is therefore plausible that MD activity in response to difficulty could also dynamically adapt according to the range of difficulty levels within a given task context, especially if it is involved in signaling difficulty.

On the other hand, an argument can also be made against context-dependent coding of difficulty. While humans can easily represent near-unlimited bounds of value (i.e., $0.01, $10, $10000, etc.), and this large range may promote contextual adaptation in terms of neural coding, the range of difficulty of information processing we can handle seems to be rather limited (Marois & Ivanoff, 2005). Capacity limits in cognitive processing include the number of items we can attend to (Chun & Marois, 2002) and hold in working memory (e.g., Miller, 1956), processing bottlenecks that hinder parallel task execution (Pashler, 1994), and the speed with which information can be encoded into working memory (Dux & Marois, 2009; Zivoni & Lamy, 2022). Various authors have linked these capacity limitations to the MD network (Marois and Ivanoff 2005; Watanabe and Funahashi 2014; Duncan et al. 2020) and, corresponding to the limited range of cognitive processing, the MD network’s capacity to adapt its response to a wide range of difficulty levels may also be limited. Specifically, several studies have found that, rather than showing a monotonic increase with task difficulty, MD activity displayed an inverted U-shape response (Callicott et al. 1999; Linden et al. 2003) or a plateau after a certain difficulty level (Todd and Marois 2004; Marois and Ivanoff 2005; Mitchell and Cusack 2008), especially when performance improvement becomes impossible even with maximal attention. Thus, activity in the MD network may reflect the investment of attentional resources, rather than objective or even subjective difficulty per se (Han and Marois 2013; Wen et al. 2018). If MD activity reflects resource investment, then this activity should increase whenever demand increases, but it should be unaltered by the difficulty of other tasks within its shared context.

The current experiment was designed to tease apart these two possibilities by creating two difficulty contexts (easy and hard). Within each context we manipulated difficulty over three levels (low, medium, high) with basic arithmetic problems. Crucially, the highest difficulty level within the easy context was matched with the lowest level in the hard context. If MD activity were context-dependent, we would expect the MD network to adapt its range of activation according to relative task difficulty within each context. Accordingly, the MD network would show a different neural response to the matched difficulty conditions across contexts, with greater activity for the high difficulty level in the easy context than for the low difficulty level in the hard context. It is also possible for MD activity to show sensitive to both context-dependent (relative) difficulty as well as context-independent (absolute) difficulty, and in this case, we would expect an additive mix in the activation response. As another test of context-dependence, we examined whether MD activity during a given trial is sensitive to the difficulty level of the previous trial. Complementing these univariate analyses, we explored representational distances of difficulty in the MD network with RSA. Finally, a whole-brain analyses was conducted to identify additional regions that may differentially represent context-dependent and context-independent responses (Grabenhorst and Rolls 2009) to difficulty.

## Materials and Methods

### Participants

The study design was based on Nieuwenhuis et al. (2005), who observed significant range adaptation effects in reward processing with a sample size of 14 individuals. We aimed to increase power via a larger sample and targeted a minimum sample size of 24, which is typical for functional imaging studies relating to the MD network (e.g., Shashidhara et al. 2019). 25 participants (9 males, 16 females; ages 18-35, mean = 25.01, SD = 4.11) were included in the analysis of this experiment. Two additional participants were excluded due to low accuracy and excessive motion during the scans (mean accuracy < 70% and/or motion > 4 mm on one or more runs). All participants were neurologically healthy with normal or corrected-to-normal vision. Procedures were conducted in accordance with ethical approval obtained from the Duke University Health System Institutional Review Committee, and participants provided written, informed consent before the start of the experiment.

### Stimuli and task procedures

The experimental design was modeled closely on Nieuwenhuis et al. (2005), but instead of reward, we manipulated task difficulty. The study consisted of an online practice session and a main experimental session in the scanner. The practice session was performed on participants’ own computers within a week before the main experiment. During both sessions, participants were told that on each trial, they would be shown three doors from either a blue set or red set. They were informed that (a) one set of doors contains more difficult problems than the other set and (b) within each set of doors, there would be three levels of difficulty (low, medium, and high), and each door is associated with one level of difficulty. Thus, the two sets of doors defined the two difficulty contexts in the experiment. Additionally, participants were told that before the beginning of each trial, the position of the doors within the presented set would be shuffled, and they were given an animation demo of the doors being shuffled during the instructions to incentivize participants to choose different locations. Behind the “easy” set of doors, the math problems could be addition of (1: low difficulty) two single digits, with the constraint of the sum not exceeding 10 (e.g., 3 + 1), (2: medium difficulty) a single digit and a double digit, with the ones position requiring a carryover (e.g., 94 + 8), or (3: high difficulty) two double digits, with at least one carryover (e.g., 26 + 57). Behind the “hard” set of doors, the math problems could be (1: low difficulty) two double digits (e.g., 19 + 42), (2: medium difficulty) a double digit and a triple digit (e.g., 925 + 86), or (3: high difficulty) two triple digits (e.g., 718 + 503), all requiring at least one carryover. Thus, the high difficulty condition in the easy set was equivalent to the low difficulty condition in the hard set. The assignment of the red and blue doors to easy versus difficult sets was counterbalanced across participants.

Figure 1 illustrates the structure of the experimental paradigm. On each trial, participants were first shown three doors from one set (i.e., the contextual cue) on the screen and had up to 2.5 s to press one out of three buttons (the “8”, “9”, and “0” keys on their keyboard during the online practice and the first three buttons of the right-hand button box in the scanner) to select the left, center, or right door. Participants were encouraged to respond to every trial, however, if the participant did not respond within the time limit, the computer would choose a door for them. As soon as a door was chosen, the word “Chosen!” would be displayed along with arrows indicating the selected door for the remainder of the 2.5 s and an additional 1 s. This was then followed by a 1.5 s fixation cross. Next, participants were presented with a screen displaying a math problem in the center with three choices below. They were given up to 6 s to select the correct answer, using the same three buttons. After an answer was chosen, participants would be shown a fixation cross for the remainder of the duration and an additional 3 s before moving on to the next trial.

**Figure 1.**
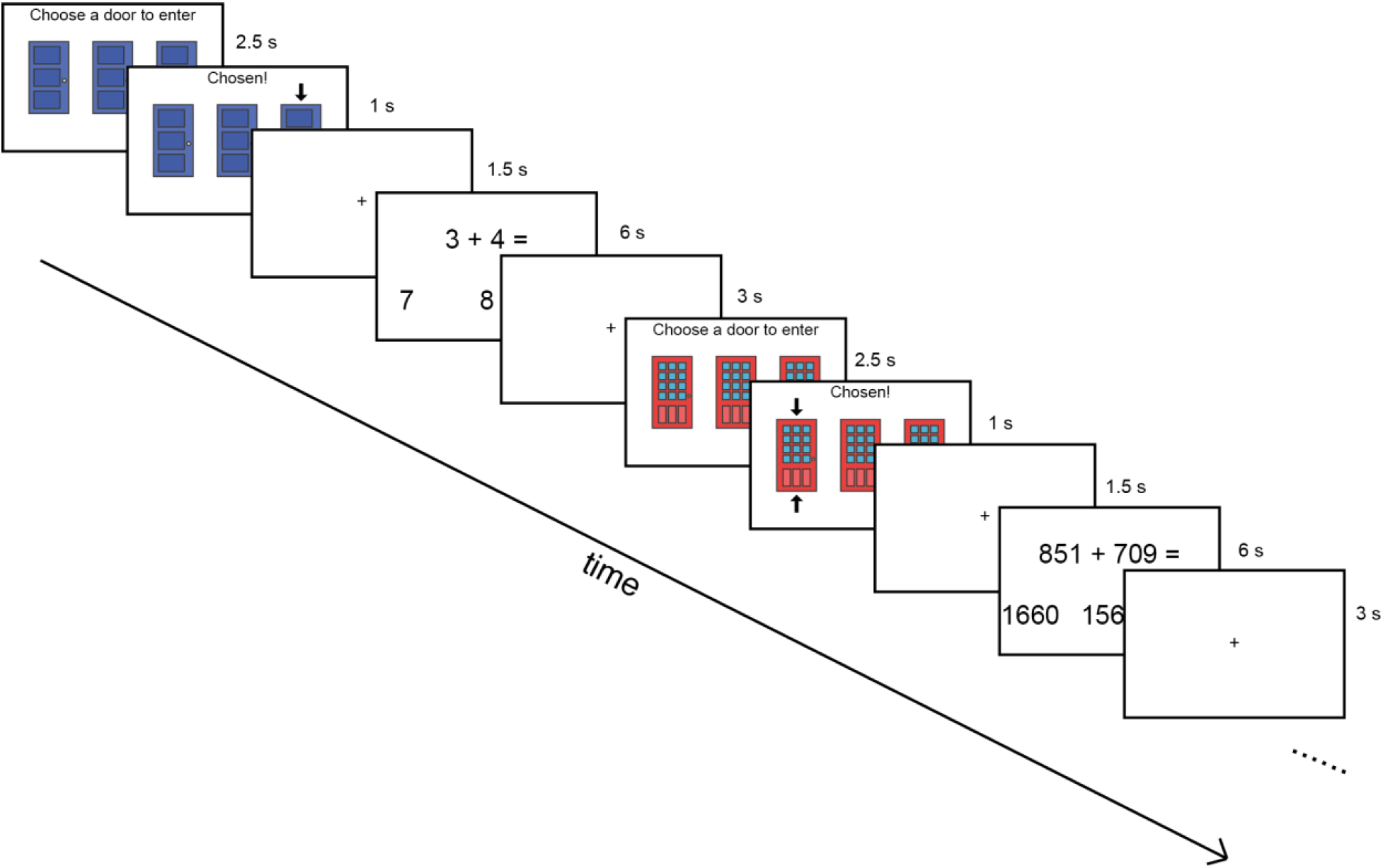
Illustration of experimental paradigm, showing two trials. On each trial, participants were given a contextual cue in the form of a set of three colored doors (blue or red, associated with easy or hard problems, counterbalanced across participants), and were asked to choose one door to enter. After selecting a door, they were given a math problem from one of the three difficulty levels associated with that colored set and were asked to select the correct answer among three choices. See text for the timing of each display.

The timing of the displays was the same for the online practice and the scanning sessions, except for that in the practice session, participants were additionally given feedback after each answer to a math problem (either “Too slow! Press the spacebar to continue”, or “correct” or “incorrect” for 500 ms). To promote learning of the associations between the colored doors and difficulty context during the online practice session, we used a blocked design (Flesch et al. 2018), where participants were given the same set of doors for 9 consecutive trials in alternating blocks. Participants performed a total of 54 practice trials in total and experienced each set of doors and difficulty level both equal number of times.

In the main experiment, participants performed 5 scanning runs. Each run had a total of 37 trials, with the first trial being a dummy trial. The two possible sets of doors and the six possible difficulty conditions (five levels) were varied on a trial-to-trial basis and occurred equally often across the experiment. The probability of switching from one condition to any other was equated, such that all possible transitions occurred equally often across the experiment. Although participants were told that different doors led to different math problems, in reality, the order of difficulty conditions was pre-determined and the sequence of difficulty conditions was unaffected by their response choices. Participants were not given feedback after each trial, but were shown an overall accuracy score after the end of each run.

### fMRI data acquisition

Scanning took place in a 3T Siemens Prisma scanner at the Center for Advanced Magnetic Resonance Development at Duke University Hospital. Functional images were acquired using a multiband gradient-echo echoplanar imaging (EPI) pulse sequence (TR = 2000 ms, TE = 30 ms, flip angle = 90°, 128 × 128 matrices, slice thickness = 2 mm, no gap, voxel size 2 × 2 × 2 mm, 69 axial slices covering the entire brain, three slices acquired at once). The first five volumes served as dummy scans and were discarded to avoid T1 equilibrium effects. A reverse phase encoding image was collected at the end of the experiment. High-resolution anatomical T1-weighted images were acquired for each participant using a 3D MPRAGE sequence (192 axial slices, TR = 2250 ms, TI = 900 ms, TE = 3.12 ms, flip angle = 9°, field of view = 256 × 256 mm, 1-mm isotropic resolution).

### Preprocessing

Preprocessing was performed using fMRIPrep 20.2.3 (Esteban et al., 2018; RRID:SCR_016216), which is based on Nipype 1.6.1 (Gorgolewski et al., 2011; RRID:SCR_002502).

#### Anatomical data preprocessing

The T1-weighted (T1w) image was corrected for intensity non-uniformity (INU) with N4BiasFieldCorrection (Tustison et al. 2010), distributed with ANTs 2.3.3 (Avants, Epstein, Grossman, & Gee, 2008; RRID:SCR_004757), and used as T1w-reference throughout the workflow. The T1w-reference was then skull-stripped with a Nipype implementation of the antsBrainExtraction.sh workflow (from ANTs), using OASIS30ANTs as target template. Brain tissue segmentation of cerebrospinal fluid (CSF), white-matter (WM) and gray-matter (GM) was performed on the brain-extracted T1w using fast (FSL 5.0.9, RRID:SCR_002823; Zhang, Brady, & Smith, 2001). Brain surfaces were reconstructed using recon-all (FreeSurfer 6.0.1, RRID:SCR_001847; Dale, Fischl, & Sereno, 1999), and the brain mask estimated previously was refined with a custom variation of the method to reconcile ANTs-derived and FreeSurfer-derived segmentations of the cortical gray-matter of Mindboggle (RRID:SCR_002438; Klein et al., 2017).

Volume-based spatial normalization to one standard space (MNI152NLin2009cAsym) was performed through nonlinear registration with antsRegistration (ANTs 2.3.3), using brain-extracted versions of both T1w reference and the T1w template. The following template was selected for spatial normalization: ICBM 152 Nonlinear Asymmetrical template version 2009c (Fonov, Evans, McKinstry, Almli, & Collins, 2009; RRID:SCR_008796; TemplateFlow ID: MNI152NLin2009cAsym).

#### Functional data preprocessing

For each of the 5 BOLD runs per subject, the following preprocessing was performed. First, a reference volume and its skull-stripped version were generated using a custom methodology of fMRIPrep. A B0-nonuniformity map (or fieldmap) was estimated based on two (or more) echo-planar imaging (EPI) references with opposing phase-encoding directions, with 3dQwarp (R. W. Cox & Hyde, 1997; AFNI 20160207). Based on the estimated susceptibility distortion, a corrected EPI (echo-planar imaging) reference was calculated for a more accurate co-registration with the anatomical reference. The BOLD reference was then co-registered to the T1w reference using bbregister (FreeSurfer) which implements boundary-based registration (Greve and Fischl 2009). Co-registration was configured with six degrees of freedom. Head-motion parameters with respect to the BOLD reference (transformation matrices, and six corresponding rotation and translation parameters) are estimated before any spatiotemporal filtering using mcflirt (FSL 5.0.9; Jenkinson, Bannister, Brady, & Smith, 2002). BOLD runs were slice-time corrected using 3dTshift from AFNI 20160207 (R. W. Cox & Hyde, 1997; RRID:SCR_005927). The BOLD time-series were resampled into standard space, generating a preprocessed BOLD run in MNI152NLin2009cAsym space. All resamplings were performed with a single interpolation step by composing all the pertinent transformations (i.e. head-motion transform matrices, susceptibility distortion correction, and co-registrations to anatomical and output spaces). Gridded (volumetric) resamplings were performed using antsApplyTransforms (ANTs), configured with Lanczos interpolation to minimize the smoothing effects of other kernels (Lanczos 1964).

Prior to fMRI analyses, we removed the first 5 TRs in each run. The functional data were high-pass filtered with a cutoff at 1/128 Hz. Spatial smoothing of 10 mm full width at half maximum (FWHM) was applied for the univariate whole-brain analysis, but not for the univariate region of interest (ROI) analysis or the representation similarity analysis (RSA). For all the analyses, we controlled the false discovery rate (FDR) to correct for multiple comparisons across ROIs as well as the whole brain.

### ROIs

For the primary analysis, we focused on the MD network (see Figures 4, 6, and 7). The MD network was based on data from Fedorenko et al. (2013), selecting frontoparietal regions responsive to cognitive demands across seven diverse tasks (http://imaging.mrc-cbu.cam.ac.uk/imaging/MDsystem). MD component ROIs were separated as described in Mitchell et al. (2016), based on proximity to local maxima in the data of Fedorenko et al. (2013); and included aMFG, mMFG, pMFG, pdLFC, IPS, AI, and ACC. As MD activation tends to be largely symmetrical, left and right hemisphere ROIs were combined to form bilateral ROIs.

### Univariate activation across difficulty conditions

Statistical analyses were performed first at the individual level, using a general linear model (GLM). In our first GLM, we had a regressor for each type of math problem that was answered correctly (6 regressors: 2 contexts × 3 difficulty levels). Math problems that were answered incorrectly were removed from the analysis using a separate regressor. All math problems were modeled with the duration of each trial’s response time (or the maximum 6 s if participants failed to provide a response). This provided an estimate of each voxel’s response per-unit-time, and thus accounts for response time differences between conditions (Henson 2007; Grinband et al. 2008; Woolgar et al. 2014). This approach was used in previous univariate and multivariate fMRI studies that had considerable reaction time differences resulting from differing task difficulty across conditions (e.g., Woolgar, Hampshire, et al. 2011; Woolgar, Thompson, et al. 2011; Crittenden and Duncan 2014; Wen et al. 2018), and has been shown to provide robust statistical power, reliability, and interpretability of fMRI results (Grinband et al. 2008). We additionally had regressors for each set of contextual cues (i.e., blue and red doors; 2 regressors). The contextual cues were modeled with a fixed 3.5 s duration. Each regressor was convolved with the canonical hemodynamic response function. The six motion parameters and block means were included as regressors of no interest. The average beta estimates for individual participants were entered into a random effects group analysis.

One possibility is that the MD network shows context-independent activity (Figure 2Ai), such that MD activity would increase linearly with increased absolute difficulty of the math problems, and there would be an equivalent response to the two double digits additions (the high difficulty level in the easy set and the low difficulty level in the hard set), regardless whether it was experienced in the easy set or hard set. Another possibility is that the MD network shows context-dependent activity (Figure 2Aii), such that activation would be scaled within each set of doors. In this scenario, we would expect the low, medium, and high difficulty conditions to elicit similar neural responses across the two sets, such that the two double digits additions to show a greater neural response when experienced in the easy set (in which it is the high difficulty condition) than when it is experienced in the hard set (in which it is the low difficulty condition). Finally, it is possible that the MD network is sensitive to both relative difficulty within a context as well as absolute difficulty that is independent of context, in which case we would expect MD activation to reflect an additive mix of the former two (Figure 2Aiii). To evaluate these possibilities, for each participant, we fit a linear regression with a regressor modeling absolute difficulty ([1,2,3,3,4,5]) and a regressor modeling relative difficulty ([1,3,5,1,3,5]) to their neural response to the six types of math problems. The individual participants’ beta estimates were then entered into a random effects group analysis.

**Figure 2.**
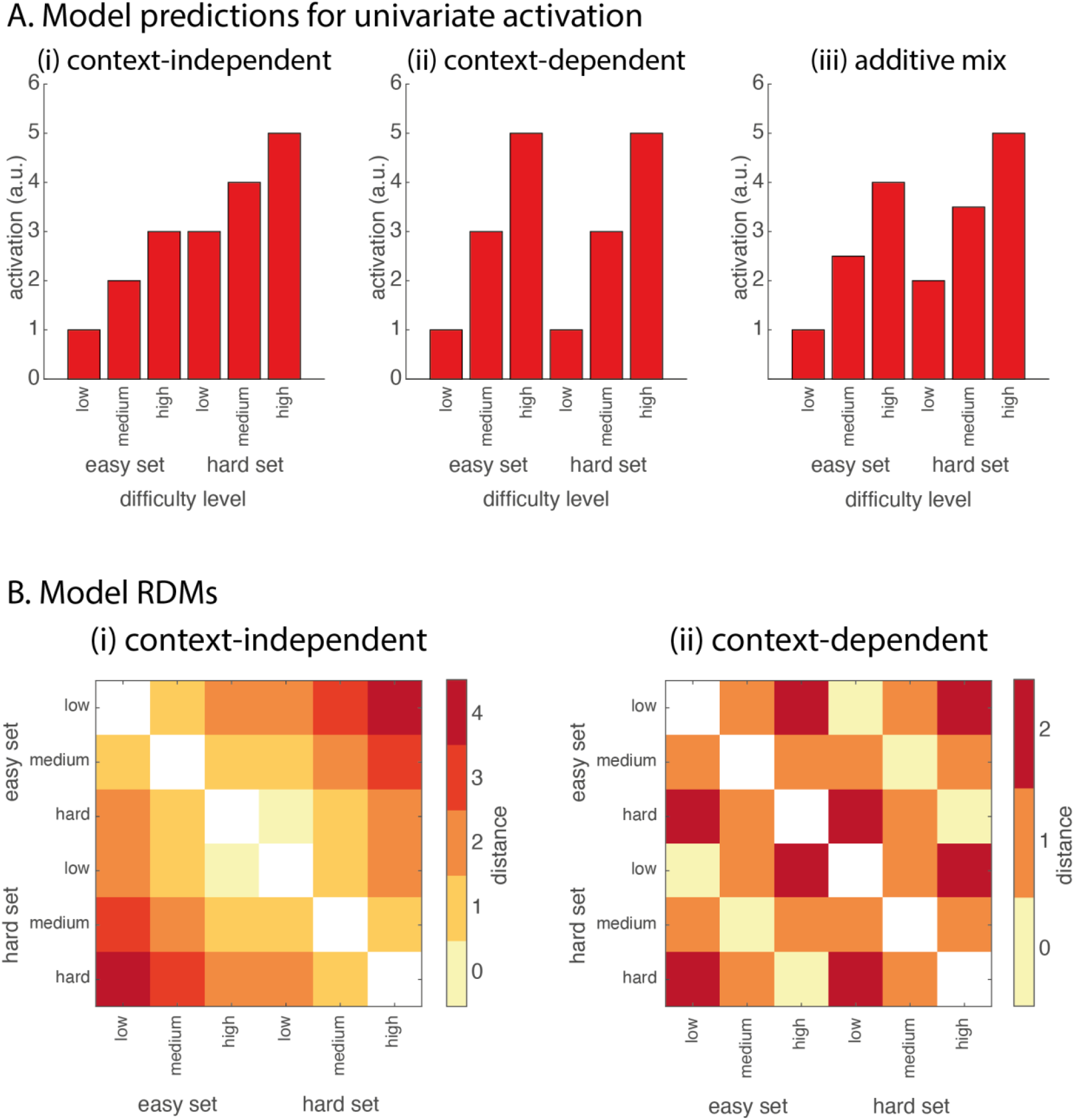
Predictions of (A) univariate activations for regions that are (i) context-independent, (ii) context-dependent, and (iii) an additive mix of the two; and (B) RDMs of (i) context-independent and (ii) context-dependent coding in difficulty processing.

### Univariate activation when switching difficulty contexts

In our second GLM, we modeled each imperative trial according to its current as well as previous difficulty context (i.e., Easy-Easy, Hard-Easy, Hard-Hard, Easy-Hard; 4 regressors). Each trial duration was modeled according to participants’ response times. We also modeled the contextual cues according to their current and previous condition (i.e., Easy-Easy, Hard-Easy, Hard-Hard, Easy-Hard; 4 regressors). The context cues were modeled with a fixed 3.5 s duration. Math problems that were answered incorrectly were removed from the analysis using a separate regressor. The first imperative trial and first cue (which did not have a previous trial to switch from) was modeled individually as a regressor of no interest (2 regressors). Each regressor was convolved with the canonical hemodynamic response function. The six motion parameters and block means were included as regressors of no interest. The average beta estimates for individual participants were entered into a random effects group analysis.

A priori, we were particularly interested in the following contrasts: (1) switching from a problem in the hard set to a problem in the easy set versus repeating a problem in the easy set, and (2) switching from the easy set to a problem in the hard set versus repeating a problem in the hard set. One demonstration of context-dependent coding would be sensitivity to previous trial experience (Akitsuki et al. 2003; Nakahara et al. 2004). We hypothesized that if a participant becomes more efficient at solving hard problems because of a previous experience with a hard trial, then we would expect decreased MD activity when switching from the hard set to the easy set (Garavan et al. 2000; Landau et al. 2004). One the other hand, we may expect increased MD activity when switching from the easy set to the hard set, as relative task difficulty would increase (Botvinick et al. 1999; Carter et al. 2000; Durston et al. 2003; Kerns et al. 2004). Thus, we would expect an interaction between switching versus repeating a set, and whether the current set is easy or hard.

### RSA analysis

We performed RSA using the linear discriminant contrast (LDC) to quantify dissimilarities between activation patterns. The analysis used the RSA toolbox (Nili et al., 2014), in conjunction with in-house software. The LDC was chosen because it is multivariate noise-normalized, potentially increasing sensitivity, and is a cross-validated measure which is distributed around zero when the true distance is zero (Walther et al., 2016). An average activity pattern for each type of math problem was obtained from the first GLM above, thus resulting in 6 patterns in total for each run. For every possible combination of two runs, and for each pair of patterns, the patterns from run 1 were projected onto a Fisher discriminant fitted for run 2, with the difference between the projected patterns providing a cross-validated estimate of a squared Mahalanobis distance. This was repeated projecting run 2 onto run 1, and we took the average as the dissimilarity measure between the two patterns. We then averaged the result from each pair of runs. All pairs of pattern dissimilarities therefore formed a symmetrical representational dissimilarity matrix (RDM) with zeros on the diagonal. This was done individually on the MD ROIs as well as in a whole-brain analysis using a 10 mm searchlight and then smoothed with a 10 mm FWHM before the group analysis. The average RDMs of MD ROIs are plotted in the Supplementary Material (Figure S4).

We constructed two model RDMs to probe for the existence of absolute, context-independent difficulty coding and relative, context-dependent difficulty coding (Figure 2B). In each RDM, each cell represents the dissimilarity between the corresponding two types of math problems. In the context-independent coding RDM, dissimilarity increases as the difference in difficulty of the math problems increases (Figure 2Bi). In the context-dependent coding RDM, the low, medium, and high difficulty levels of each set are represented with the smallest dissimilarity, and dissimilarity increases accordingly to the distance between these three levels (Figure 2Bii). We note that the two model RDMs have little correlation with each other (Spearman’s ρ= −0.04, p = 0.90).

Since the stimuli used in the different conditions also differ in visual similarity, for each participant, we constructed an additional model RDM estimating the pixel-level correlation distance between the mean math stimuli of the different conditions as they were presented on screen. The average pixel RDM across participants is illustrated in the Supplementary Material (Figure S3). The rank correlations between the pixel RDM and context-independent RDM (Spearman’s ρ= 0.36, p = 0.19) and context-dependent RDM (Spearman’s ρ= −0.11, p = 0.69) were not statistically significant. The above three model RDMs, as well as the brain RDM, were rank-transformed and entered into the following regression:

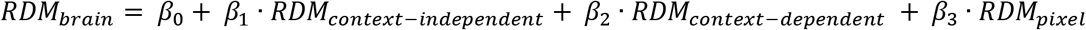

Rank-transformation was performed because we did not want to assume a linear relationship between the dissimilarities (Nili et al. 2014). Beta coefficients were estimated for each participant, and one-tailed t-tests against zero were then performed to identify ROIs or regions that showed a significant relationship between the brain RDM and model RDMs.

### Data and code sharing

All experimental stimuli and task/analysis codes are available at https://github.com/tanya-wen/Difficulty-MD-network.

## Results

### Behavioral results

As shown in Figure 3A, accuracy decreased while reaction times increased with difficulty of the math problem. Overall, accuracy decreased from a mean of 98.84%, to a mean of 72.19% from the easiest (two single digits) to the hardest (two triple digits) difficulty level. Pairwise t-tests showed no differences between the two matched levels, that is, the high difficulty condition in the easy set and the low difficulty condition in the hard set (t = 0.36, p = 0.72). There were significant differences between all other trial types (all ts > 2.69, all ps < 0.02; FDR corrected for multiple comparisons). Average median reaction time increased from 0.97 s in the easiest level to 4.12 s in the hardest level. Pairwise t-tests showed significant differences between all trial types, with the smallest difference occurring between the matched difficulty conditions where participants were slightly slower in responding to the low difficulty level in the hard set compared to the high difficulty level in the easy set (all ts > 2.64, all ps < 0.02; FDR corrected for multiple comparisons).

**Figure 3.**
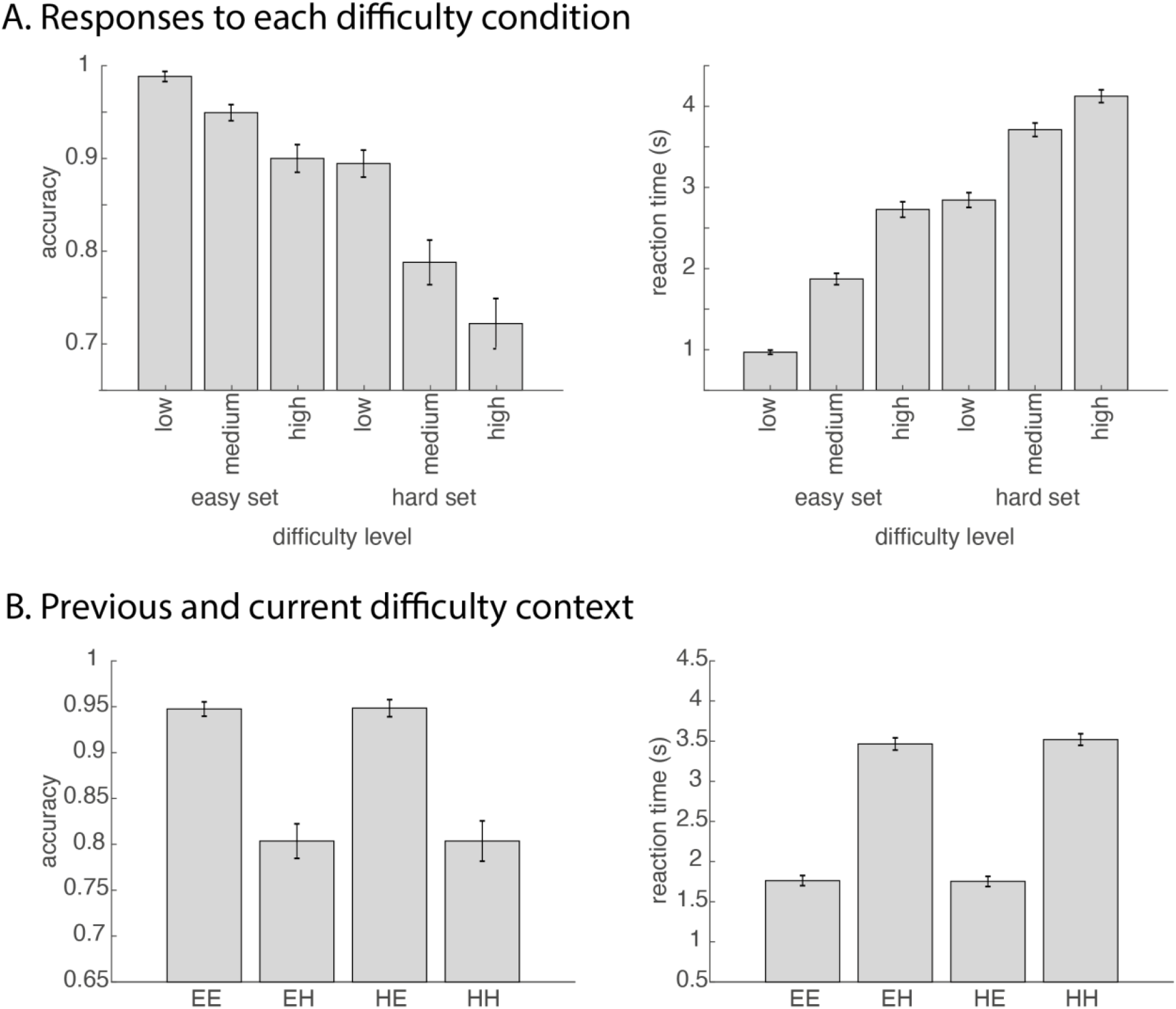
(A) Behavioral results of accuracy (left) and reaction time (right) across the six math conditions. (B) Accuracy (left) and reaction time (right) plotted as a function of the current context as well as the previously experienced context. EE: previous easy current easy; EH: previous easy current hard; HE: previous hard current easy; HH: previous hard current hard. Error bars represent standard error.

**Figure 4.**
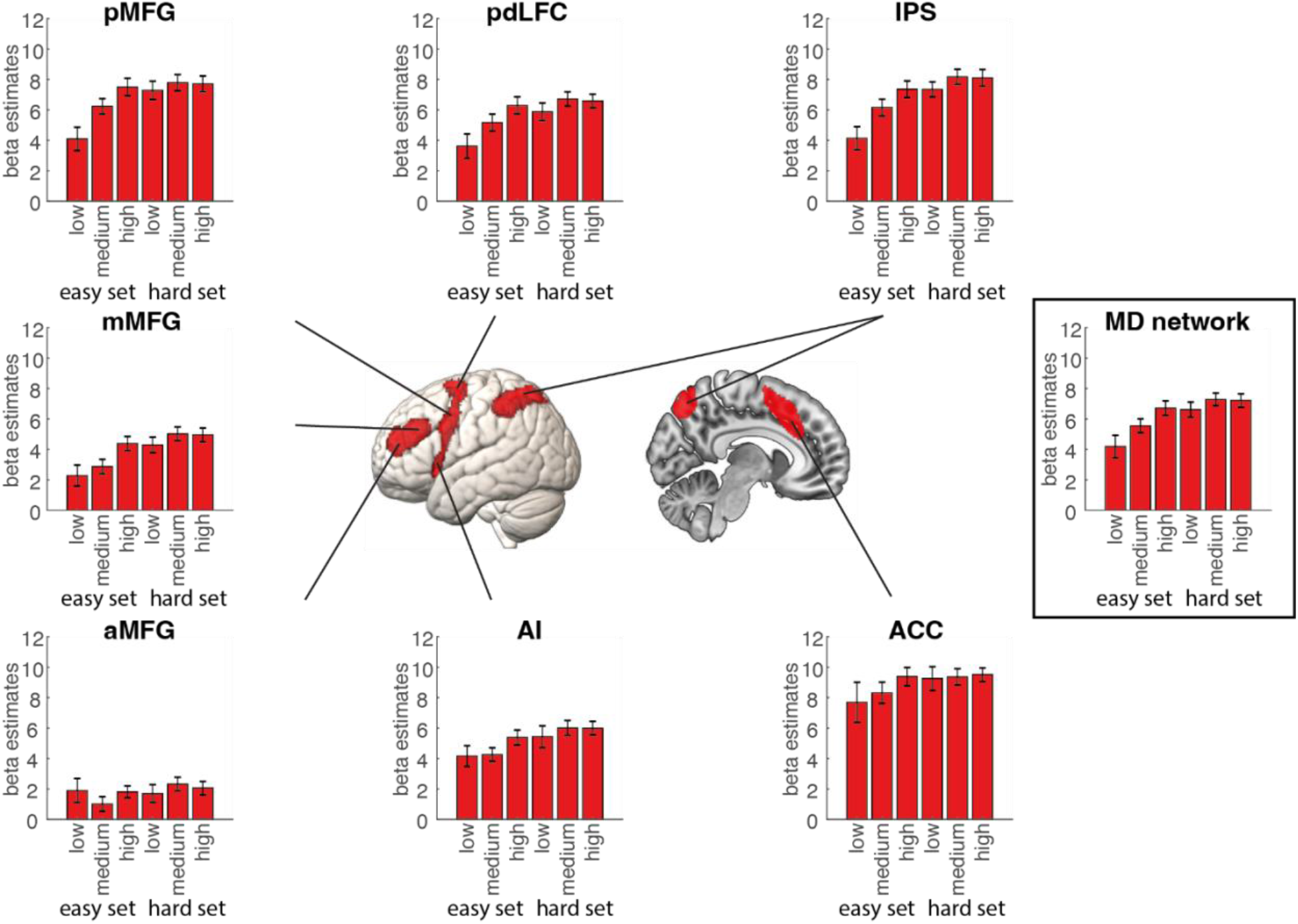
ROI results of MD regions (left) as well as the entire MD network (right). Graphs show the beta values for each math condition. Error bars represent standard error.

We next examined whether there were any behavioral signatures of context-dependence based on previously experienced difficulty contexts. First, we grouped all trials according to whether they belonged to the hard or easy set, and which set proceeded them. A two-way repeated measures ANOVA with factors of previous difficulty (easy vs. hard) × current difficulty (easy vs. hard) was performed on the accuracy data and median reaction time of correct trials, respectively (Figure 3B). For accuracy, we found a significant main effect of current difficulty (F(1,24) = 91.68, p < 0.001), which is caused by the hard set having lower accuracy than the easy set. There was no main effect of previous difficulty (F(1,24) < 0.01, p = 0.97) or previous difficulty × current difficulty interaction (F(1,24) < 0.01, p = 0.96). For reaction time, we found a significant main effect of current difficulty (F(1,24) = 1755.35, p < 0.001), with longer reaction times for the hard set. There was no main effect of previous difficulty (F(1,24) = 0.29, p = 0.59) and no previous difficulty × current difficulty interaction (F(1,24) = 1.24, p = 0.28). Thus, when analyzing sequence effects, we observed no context-dependence in accuracy or RT data.

### Univariate activation across difficulty conditions

#### ROI analysis

Average beta estimates of each difficulty level from bilateral MD regions are shown in Figure 4. We also extracted the average activation for each of the six difficulty conditions from each participant using a combined MD network ROI. We first ran a context (easy vs. hard) × difficulty level (low, medium, and high) repeated measures ANOVA to examine activity in the MD network. Results showed a significant main effect of context (F(1,24) = 21.47, p < 0.001) and a significant main effect of difficulty level (F(2,48) = 9.89, p < 0.001). There was also a context × difficulty level interaction (F(2,48) = 4.10, p = 0.02). Pairwise t-tests across the six difficulty conditions revealed several significant contrasts, with increased absolute difficulty associated with increased MD activation, although starting to plateau at the higher difficulty levels (no significant difference between the medium and high difficulty levels in the hard set (t = 0.20, p = 0.84)).

To compare across difficulty levels and ROIs, we conducted a 3-way ANOVA with factors context (easy vs. hard), difficulty level (low, medium, and high), and ROI (7 MD ROIs). This analysis showed a significant main effect of context (F(1,24) = 15.31, p < 0.001), a significant main effect of difficulty level (F(2,48) = 5.71, p < 0.01), and a significant main effect of ROI (F(6,144) = 64.81, p < 0.001). The main effect of context was driven by increased MD activity in the hard compared to easy context, and the main effect of difficulty was driven by increasing MD activity with higher difficulty levels. There was a context × ROI interaction (F(6,144) = 7.20, p < 0.001) and difficulty level × ROI interaction (F(12,288) = 5.80, p < 0.001), but no context × difficulty level interaction (F(2,48) = 2.75, p = 0.07). Finally, there was a context × difficulty level × ROI interaction (F(12,288) = 6.34, p < 0.001). Pairwise t-tests across the six difficulty conditions revealed that in most of the MD ROIs, there was a general increase and plateau in activation as difficulty increased, except for the aMFG and ACC, where the activation remained relatively similar across all six conditions. Details of corrected and uncorrected statistics per ROI are listed in the Supplementary Material (section 2.1).

Our key *a priori* hypothesis for a context-dependent system was that it would show a different neural response to the two double digit addition when experienced in the easy set than in the hard set. As these two conditions are matched in stimuli and requisite cognitive operations, any differences observed between these conditions would reflect the effect of context. Crucially, we found no difference in the MD network ROI in response to these two conditions (t = 0.31, p = 0.76). A Bayesian model comparison using JASP (JASP team, 2022) found a BF01 value of 4.54, indicating that there was 4.54 times more evidence in favor of the null model where there is no difference between the two conditions than a model where the two conditions differ, indicating substantial evidence in favor of the null hypothesis. Furthermore, none of the individual ROIs showed any significant differences between the matched difficulty conditions (all |t|s < 1.19, all ps > 0.98; FDR corrected for multiple comparisons; BF_01_ = 2.53 for the most significant ROI).

#### Whole-brain analysis

We also carried out a whole-brain analysis to examine other potential regions that may show context-independent or context-dependent activation. To do this, we fit a linear regression for the six difficulty conditions in each voxel with the regressors [1,2,3,3,4,5] and [1,2,3,1,2,3] (see Material and Methods). Results are shown in Figure 5. For the context-independent regressor, there was a significant positive association with activity throughout the MD network, largely overlapping with the ROIs, as well as in visual cortex. Significant negative association was found with activity in default mode network (DMN) regions, as these regions showed decreased activation as absolute difficulty increased. This observation is consistent with previous findings of the DMN showing decreased activity during externally oriented, cognitively demanding tasks (Raichle and Snyder 2007; Gilbert et al. 2012). There were no significant activity associations for the context-dependent regressor in either positive or negative direction.

**Figure 5.**
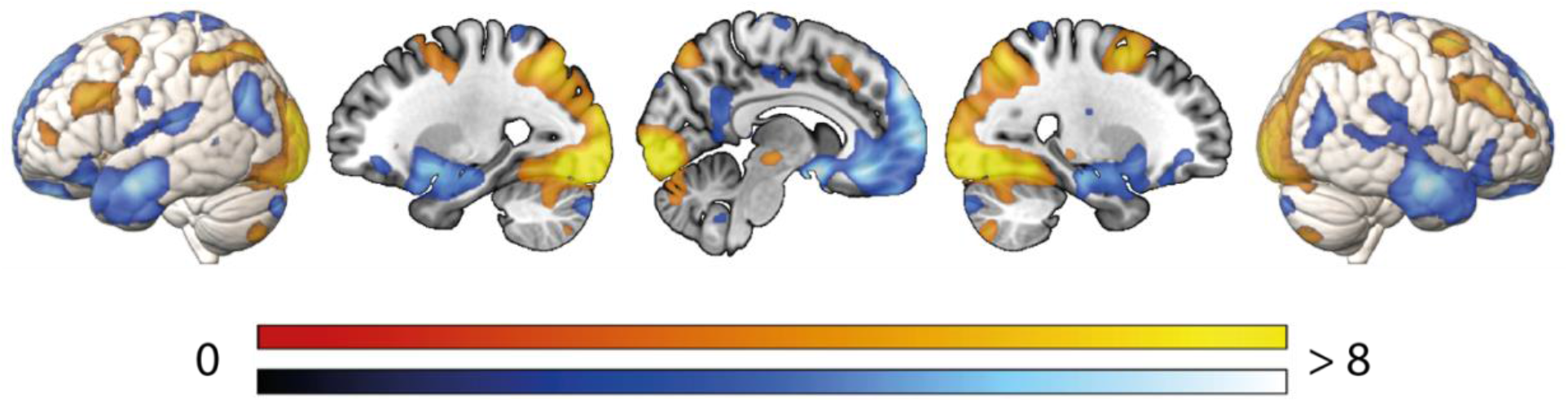
Whole-brain voxelwise regression showing significant associations to absolute, context-independent levels of difficulty. Colors indicate t values, with warm and cool scales indicating positive and negative tails, respectively. The activation maps are thresholded at FDR < 0.025 per tail.

**Figure 6.**
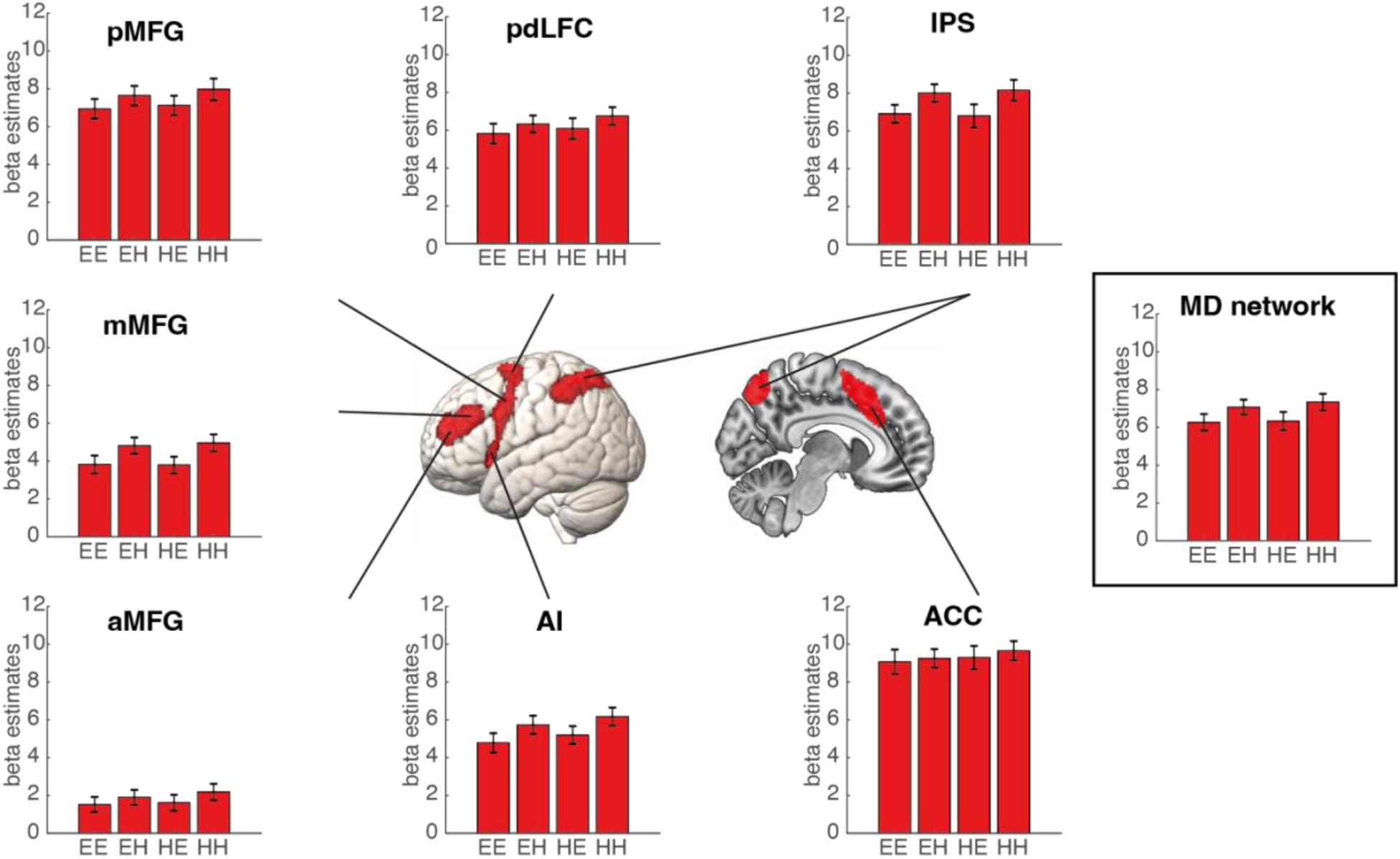
Activity in the MD network during task execution based on previous and current trial difficulty. EE: previous easy current easy; EH: previous easy current hard; HE: previous hard current easy; HH: previous hard current hard. Error bars represent standard error.

**Figure 7.**
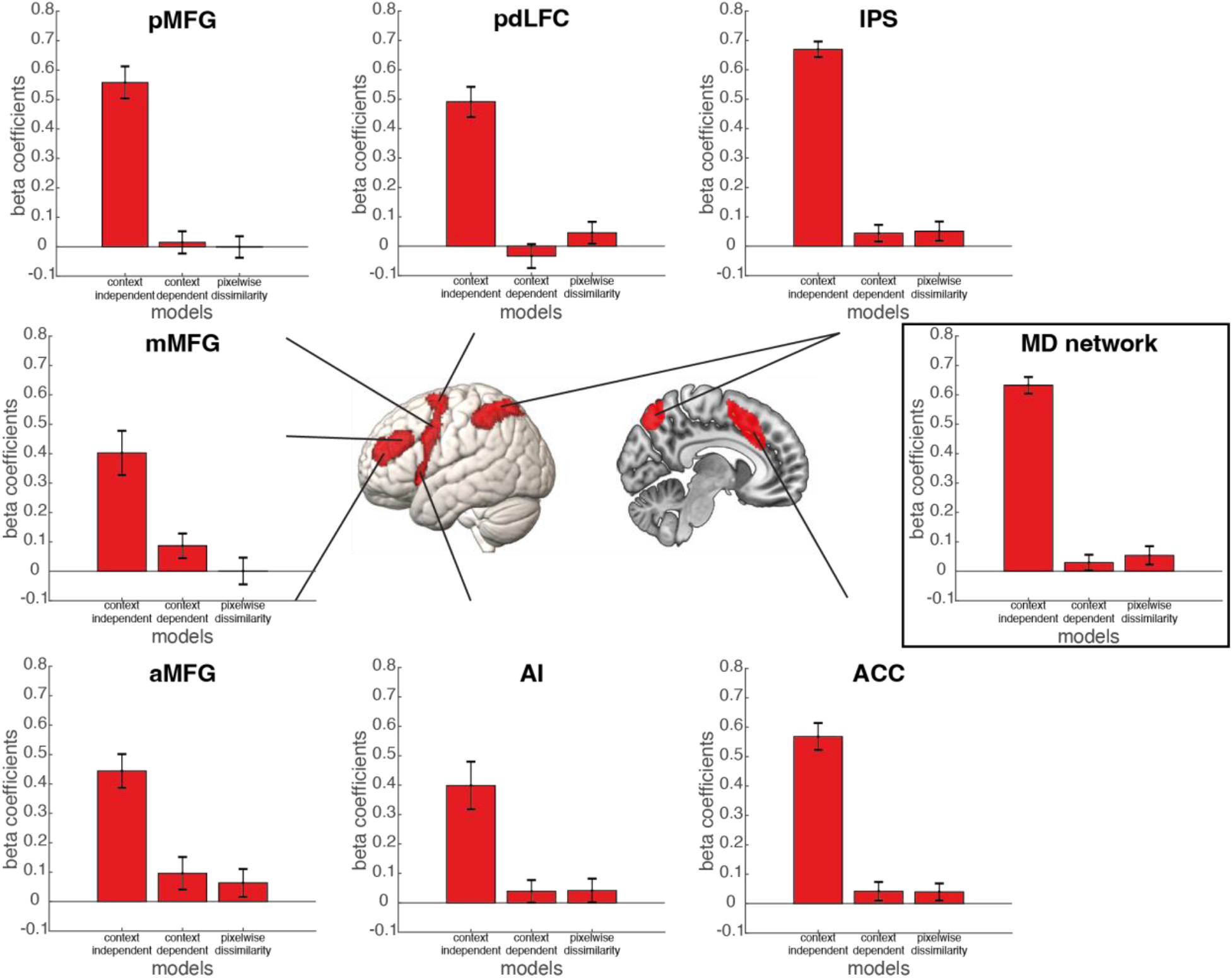
Relatedness of the model RDMs to the brain RDM for the MD ROIs (left) and the entire MD network (right). Beta coefficients were calculated by fitting the three model RDMs (context-independent, context-dependent, and pixelwise dissimilarity) to the brain RDM for each ROI after rank transformation. Error bars represent standard error.

While the DMN was not the target of our a priori hypotheses in this study, given its sensitivity to task difficulty and previous indications that it may represent task context (Crittenden et al. 2015; Smith et al. 2018; Wen, Duncan, et al. 2020; Wen, Mitchell, et al. 2020), we nevertheless performed some exploratory analysis to examine DMN region responses, as detailed in the Supplementary Material (section 5), by replicating the analyses performed the MD ROIs. These analyses correspond to the findings of the whole-brain analyses and suggest that the univariate activity of the DMN aligned with the context-independent model, showing increased deactivation with increased task difficulty (Supplementary Material, section 5.1-5.2).

Finally, as the matched difficulty conditions may be the most sensitive test for context-dependent effects, we also directly compared the high difficulty level in the easy set and the low difficulty level in the hard set, at the whole-brain level. No region showed significant differences between the two conditions at FDR < 0.05 in either direction. Thus, we obtained evidence only for context-independent coding of difficulty.

### Univariate activation when switching difficulty contexts

#### ROI analysis

Figure 6 shows the average MD response to the two sets of math problems as a function of the difficulty of the previous trial set during the processing of the math problem. We first performed a previous difficulty (easy vs. hard) × current difficulty (easy vs. hard) ANOVA on the combined MD ROI. To further examine differences between ROIs, we conducted a previous difficulty (easy vs. hard) × current (easy vs. hard) × ROI (7 MD ROIs) ANOVA.

The MD network ROI showed a significant main effect of current difficulty (F(1,24) = 17.05, p < 0.001), which was a result of higher activation when performing math problems from the hard set. There was no main effect of previous difficulty (F(1,24) = 0.51, p = 0.48, BF01 = 2.88) or previous difficulty × current difficulty interaction (F(1,24) = 0.47, p = 0.50, BF01 = 2.58). In the ANOVA with the additional factor of ROI, we found a significant main effect of current difficulty (F(1,24) = 12.85, p = 0.001) and significant a main effect of ROI (F(6,144) = 74.09, p < 0.001), but no main effect of previous difficulty (F(1,24) = 0.93, p = 0.35, BF01 = 5.69). There was a current difficulty × ROI interaction (F(6,144) = 6.62, p < 0.001), but no previous difficulty × ROI (F(6,144) = 1.13, p = 0.35, BF01 = 382.39), previous difficulty × current difficulty (F(6,144) = 0.35, p = 0.56, BF01 = 11.04), or previous difficulty × current difficulty × ROI interaction (F(6,144) = 0.09, p > 0.99, BF01 = 1.59 × 10^6^). Separate previous difficulty × current difficulty ANOVAs on each of the MD ROIs showed that similar to the previous GLM, all ROIs, except for the aMFG and ACC (both Fs(1,24) < 3.52, p > 0.07) showed a main effect of difficulty (all Fs(1,24) > 6.79, all ps < 0.02). The main effects of difficulty were driven by higher activation during execution of hard versus easy math problems. Details of corrected and uncorrected statistics per ROI are listed in the Supplementary Material (section 2.2).

We additionally examined whether the MD network was sensitive to previous trial difficulty during the context cue, as presented in the Supplementary Material (section 1.1). Results showed some regions within the MD network, including the ACC, pdLFC, and pMFG displayed increased activation when the current context cue signaled an upcoming hard math problem. Moreover, the ACC and pdLFC may be sensitive to the previous difficulty context during the cue, as indicated by increased activation to the context cue if the previous trial came from the easy set (Supplementary Material, section 1.1). Exploratory analysis of these contrasts in the DMN are presented in the Supplementary Material (section 5.3), which suggested that the DMN may be sensitive to difficulty context during cue processing; however, during task execution, we were only able to identify DMN sensitivity towards the current and not previous difficulty context.

In summary, these results suggest that some regions within the MD network may be sensitive to difficulty context during cue processing; however, during task execution, the MD network seemed only sensitive to the current difficulty level, unaffected by the context of task difficulty on the previous trial.

#### Whole-brain analysis

We examined responses including the main effect of previous difficulty, main effect of current difficulty, switching to an easy set versus repeating an easy set, and switching to a hard set versus repeating a hard set at the whole-brain level to examine possible effects outside of the MD network. These contrasts correspond to the components of a previous difficulty (easy vs. hard) × current difficulty (easy vs. hard) ANOVA. Results from this whole-brain analysis are presented in the Supplementary Material (Figure S2).

During the math problem, no brain region was found to show a main effect of previous difficulty. There was significant activation throughout the MD network, as well as in the visual cortex, for hard versus easy problems. Significant activation was found in DMN for easy versus hard problems. No regions were identified when switching to an easy set versus repeating an easy set, nor its reverse contrast. Finally, no regions were identified when switching to a hard set versus repeating a hard set, although the thalamus and claustrum were more activated when repeating a hard set compared to switching from an easy to hard set.

Analyses performed during the contextual cue are also presented in the Supplementary Material (Figure S2).

In summary, we identified several regions both within and outside the MD network that show context-dependence during the context cue, such that previously experienced task difficulty influenced activation levels in these regions during the cue. However, no brain region showing context-dependence during task execution was identified. Instead, we found that the MD network was more active when solving math problems from the hard compared to easy set. These results suggest MD activity during task execution is context-independent.

### RSA results

#### ROI analysis

We first performed RSA on the MD ROIs, fitting the rank-transformed brain RDM with the rank-transformed context-independent, context-dependent, and pixel RDMs using a linear regression on each participant. Results are shown in Figure 7. In all MD ROIs, the beta coefficients of the context-independent model RDM were significantly greater than zero, indicating a relationship with the brain RDMs (all ts > 4.91, all ps < 0.001; FDR corrected for multiple comparisons) and provided a significantly better fit than the context-dependent model RDM (all ts > 3.74, all ps < 0.01; FDR corrected), the coefficients of which were not significantly greater than zero (all |t|s < 2.09, all ps > 0.15; FDR corrected; BF01 = 0.39 for the most significant ROI). Finally, the beta coefficients for the pixel RDM also did not differ significantly from zero (all |t|s < 1.56, all ps > 0.20; FDR corrected; BF01 = 0.88 for the most significant ROI). For the combined MD network ROI, the same pattern was observed. The beta coefficients of the context-independent model RDM were significantly greater than zero (t = 22.59, p < 0.001) and had a significantly better fit than the context-dependent model RDM (t = 14.75, p < 0.001), which was not significantly greater than zero (t = 1.10, p = 0.14, BF01 = 1.62). The beta coefficients for the pixel RDM were also not significantly greater than zero (t = 1.70, p > 0.05, BF01 = 0.72). Details of corrected and uncorrected statistics per ROI are listed in the Supplementary Material (section 2.3).

The above analyses were also performed on the DMN in the Supplementary Material (section 5.4). Results suggested that in all DMN ROIs, the beta coefficients of the context-independent model RDM were significantly greater than zero. Additionally, in some DMN ROIs, including the dMPFC, LTC, TPJ, Rsp, as well as the entire DMN as a single ROI, we found the coefficients of the context-dependent model RDM were significantly above zero. These results suggest that the DMN may exhibit both context-independent and context-dependent coding of difficulty.

Finally, for the MD network, we also independently compared the LDC values of the matched difficulty conditions, arguably providing the most sensitive test for context-dependent effects. We did this by testing whether the distance between the high difficulty level in the easy set and the low difficulty level in the hard set was significantly greater than zero across subjects. Across all MD ROIs, we did not find any ROI showing this effect (all ts < 1.06, all ps < 0.97, BF01 = 1.72 for the most significant ROI). Details of corrected and uncorrected statistics per ROI are listed in the Supplementary Material (section 2.4). The distance between the matched conditions in the MD network ROI was also not significantly greater than zero (t = −1.23, p = 0.89, BF01 = 9.64). Thus, we were unable to find evidence for context-dependent coding in the MD network.

#### Whole-brain searchlight

To explore the effects of context-independent and context-dependent coding effects outside the a priori MD ROIs, we carried out an RSA searchlight analysis. Results are shown in Figure 8. Context-independent representation of difficulty was significant across large areas of the brain, although it was strongest in the visual cortex and local peaks in MD and DMN regions (Figure 8A). This aligns with our univariate results of strong context-independent activation in these regions. We note that the large swathes of activation may also partially be due to spatial smoothing from the 10 mm searchlight. We observed several regions showing context-dependent coding, including the inferior parietal lobule, paracentral lobule, posterior insula, and large areas of the visual cortex (Figure 8B). Since these regions are observed to be correlated with both context-independent and context-dependent RDM models (which are largely orthogonal), we suggest that they show sensitivity to both absolute difficulty as well as the contextual difficulty in which the task was presented. We did not find any significant regions that corresponded to the pixel RDM, thus ruling out that any of the above findings reflect merely pixel-by-pixel feature similarity.

**Figure 8.**
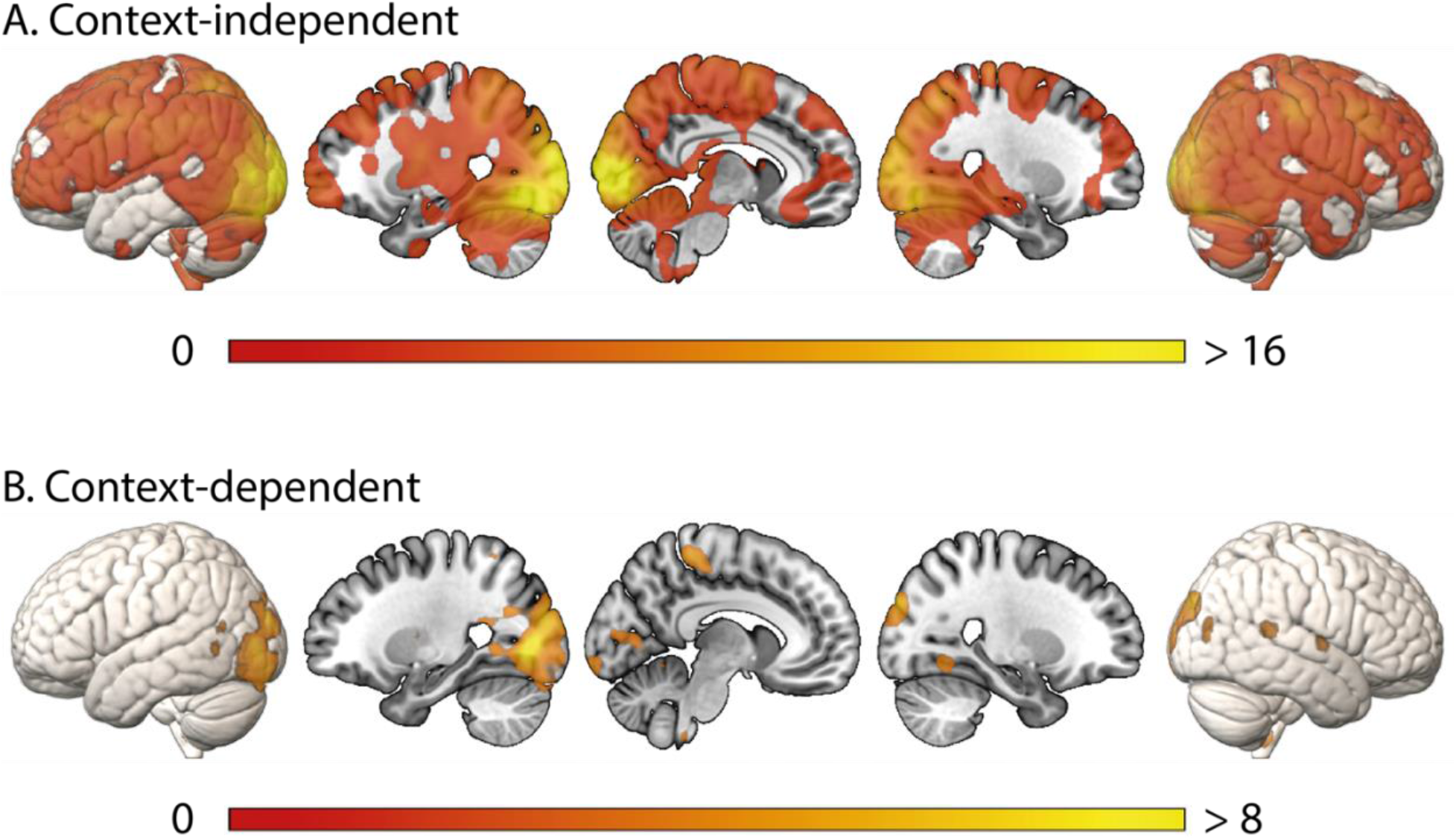
(A) Context-independent and (B) context-dependent coding of difficulty across the whole brain, calculated using local spherical searchlights, and thresholded at FDR < 0.05.

## Discussion

The present study examined for the first time whether activity in the MD network is responsive to task difficulty in a context-dependent or context-independent manner. Univariate activations as well as RSA analysis suggested that the response of the MD network to difficulty is context-independent, such that activations increased with the absolute difficulty of the task and representational dissimilarity increased with the difference of difficulty between levels, rather than being re-scaled between contexts comprising easier or harder trials. Accordingly, identical difficulty levels across the two contexts elicited equivalent MD activity, even though they represented the highest difficulty level in the easy context and the lowest difficulty level in the hard context. These results are inconsistent with context-dependent effects observed in value and sensory processing, but are consistent with the notion that strong MD activation during more difficult tasks reflects the increased demand in integrating the components of cognitive operations to solve the task at hand (Duncan 2013; Duncan et al. 2020).

In everyday life, perceived difficulty may sometimes seem relative to previous experiences. For example, if one completes mock exam that is harder than the actual test, the test will feel easy; on the other hand, if one fails to adequately prepare, then the test would feel hard (Bjork et al. 2013; Carpenter et al. 2020). We found some effect of the previous trial on the current difficulty context during the cue presentation in the ACC and pdLFC of the MD network, as well as in the DMN. These results are consistent with previous studies on sequence effects in the ACC and regions in the prefrontal cortex in the conflict-control literature, where an incongruent trial following a congruent trial elicits more activation than repeating an incongruent trial (Botvinick et al. 1999; Carter et al. 2000; Durston et al. 2003; Kerns et al. 2004). Yet, during the actual execution of the task, we were not able to identify an effect of the previous context in any of the MD ROIs in the present study, which again suggests MD is not sensitive to relative difficulty.

Why would brain activity related to cognitive demand be mostly context-independent when value-based and sensory processing commonly display range adaptation effects? One common explanation for context-dependent coding is that neural computation is costly and maximum firing rates are limited, so an efficient neural code should adapt to the range of possible values within the present context (Padoa-Schioppa 2009; Louie and Glimcher 2012; Cox and Kable 2014; Glimcher 2014). This allows humans to represent and compare seemingly unlimited ranges of values, from fractions to trillions, without much increase in effort or cost to performance. However, our capacity for high-level cognitive operations is inherently limited, and when cognitive processes become overloaded, there is degradation in performance (Norman and Bobrow 1975). In other words, in cognitive tasks, such as solving an arithmetic problem, performance is a function of the amount of cognitive resources available, and thus hits a natural limit (Kahneman 1973; Norman and Bobrow 1975; Marois and Ivanoff 2005). As previous studies have shown, MD network activity may reflect the degree to which performance can be improved by increasing attentional investment (Han and Marois 2013; Wen et al. 2018). If this were the case, then MD activity should be context-independent, as other trials should not affect the number of cognitive processes required for any given trial. Another way of looking at this is that, unlike the vast range of possible rewards values or sensory stimulation, the range of cognitive processing demand that can be handled by the brain is so limited that contextual adaptation for range coverage is unnecessary.

We note that our experiment is an event-related design, such that difficulty changed trial by trial. It therefore remains possible that MD activity might adapt to different difficulty ranges after longer exposure. Several studies have shown that task-related brain activity in MD regions may decrease after practice on a task (Garavan et al. 2000; Jansma et al. 2001; Milham et al. 2003; Landau et al. 2004), presumably reflecting increased neural efficiency, with fewer neural resources required to achieve the same level of performance. Thus, future studies may test whether MD activity would be sensitive to context in a blocked design (Bavard et al. 2021) or with separate groups of subjects experiencing different ranges of difficulty levels. Having said that, it should be noted that the coding of reward outcomes in a near-identical event-related design was found to be context-dependent (Nieuwenhuis et al., 2005). We can therefore conclude that trial-by-trial changes in context do not generally pre-empt context adaptation effects, and that the processing of task difficulty seems to fundamentally differ from the processing of reward outcomes.

Using whole-brain analysis, we furthermore explored whether there were regions outside the MD network that may show context sensitivity to task difficulty. We did not find any regions that activated in accordance with the univariate predictions of a context-dependent model. However, our RSA searchlight uncovered several regions, most notably the inferior parietal lobule, paracentral lobule, posterior insula, and large areas of the visual cortex whose activity patterns were associated with context-dependent coding. Additional exploratory analysis on the DMN revealed several ROIs, as well as the DMN as a combined ROI, to show context-dependent coding, consistent with previous studies showing context representation in the DMN (Crittenden et al. 2015; Smith et al. 2018; Wen, Duncan, et al. 2020; Wen, Mitchell, et al. 2020). It has been proposed that the posterior parietal cortex encodes abstract relational information among stimuli and the structure of the environment (Summerfield et al. 2019). Furthermore, regions in the medial temporal lobe have been identified to map conceptual space of a task (Theves et al. 2020). These results are in line with the notion that the brain is capable to matching relational knowledge of levels (low, medium, and high) across different contexts (Sheahan et al. 2021).

Previous studies have shown that mental effort is costly and has negative utility (Kool and Botvinick 2018), and when given the choice, participants typically choose tasks or contexts with low compared to high cognitive demand (Kool et al. 2010). It would therefore seem plausible that some reward-sensitive (cost-sensitive) brain regions should track task difficulty or cognitive effort. Our univariate analysis found that MD network regions showed increased activation and DMN regions showed decreased activation as the absolute difficulty of the task increased, regardless of context. Meta-analyses of the valuation network have documented that it has substantial overlap with several regions in the MD and DMN networks, including the ACC and AI in the MD network, and vmPFC and PCC in the DMN network (Levy and Glimcher 2012; Bartra et al. 2013), so these regions may have been involved in tracking effort cost in the current paradigm. However, it is difficult to evaluate where coding of effort costs is located in the brain without fully crossing reward and effort variables to decorrelate reward and difficulty (Westbrook et al. 2019). As our study did not manipulate reward, we cannot know with certainty the relationship between the two variables in the present data. For example, it is possible that correctly solving a hard problem could be more rewarding (e.g., less boring, a bigger accomplishment) than solving an easy problem (Wu et al. 2021). We also acknowledge that failure to detect a context-dependent effect under the null hypothesis significance testing does not necessary indicate lack of context coding.

Context-dependent representation in the brain seem ubiquitous in many domains, including sensory processing (Carandini and Heeger 2011; Cheadle et al. 2014), temporal perception (Walker et al. 1981; Murai et al. 2016), reward (Cox and Kable 2014; Bavard et al. 2018), and value (Sheahan et al. 2021). However, our study showed that the response of the MD network to task difficulty may be an exception to the norm. While context-dependent coding can be useful to compare values within the currently relevant context, it often leads to irrational choices, such as picking a suboptimal option under certain manipulations (Kahneman and Tversky 1979; Tversky and Simonson 1993; Chung et al. 2017; Bavard et al. 2021). Absolute coding is important for consistent and rational choices (Lee et al. 2007; Padoa-Schioppa and Assad 2007; Grabenhorst and Rolls 2009). In one study, Chung et al. (2017) found that stronger functional connectivity between frontal and reward regions when participants successfully overrode the decoy effect and made unbiased choices. Accordingly, it is possible that context-dependent coding in some areas, such as sensory and reward regions, combined with context-independent coding in MD regions, together contribute to adaptive human behavior.

## Acknowledgements

We thank Yuxi Candice Wang for assisting with fMRI data collection. We would also like to thank Daniel Mitchell for helpful discussions. This work was supported by the National Institute of Mental Health of the National Institute of Health (R01 MH116967-01A1 to T.E.).

