## Supplementary Material for "Context-independent scaling of neural responses to task difficulty in the multiple-demand network"

Table of Contents

1. [Univariate activation when switching difficulty contexts	3](#_Toc118968110)

1.1. [ROI analysis	3](#_Toc118968111)

1.2. [Whole-brain analysis	4](#_Toc118968112)

2. [Corrected and uncorrected statistics per ROI	7](#_Toc118968113)

2.1. [Univariate activation across difficulty conditions	7](#_Toc118968114)

2.2. [Univariate activation when switching difficulty contexts	11](#_Toc118968115)

2.2.1. [Context Cue	11](#_Toc118968116)

2.2.2. [Task Execution	12](#_Toc118968117)

2.3. [RSA regression	13](#_Toc118968118)

2.4. [LDC distances for matched conditions	14](#_Toc118968119)

3. [Pixelwise dissimilarity	15](#_Toc118968120)

4. [Empirical RDMs of MD regions	16](#_Toc118968121)

5. [Analysis of the DMN	17](#_Toc118968122)

5.1. [ROIs	17](#_Toc118968123)

5.2. [Univariate activation across difficulty conditions	17](#_Toc118968124)

5.3. [Univariate activation when switching difficulty contexts	19](#_Toc118968125)

5.3.1. [Context cue	20](#_Toc118968126)

5.3.2. [Task execution	21](#_Toc118968127)

5.4. [RSA results	23](#_Toc118968128)

5.5. [Empirical RDMs of DMN regions	25](#_Toc118968129)

6. [References	26](#_Toc118968130)

### 1. Univariate activation when switching difficulty contexts

#### 1.1. ROI analysis

For activity during the context cue presentation (Figure S1), the MD network ROI showed no main effect of previous difficulty (F(1,24) = 1.32, p = 0.26), no main effect of current difficulty (F(1,24) = 4.01, p = 0.06), and no previous difficulty × current difficulty interaction (F(1,24) = 2.37, p = 0.14). Adding the 7 ROIs as an additional factor of interest, the ANOVA results showed a main effect of current difficulty (F(1,24) = 5.76, p = 0.02), a main effect of ROI (F(6,144) = 49.94, p < 0.001), but no main effect of previous difficulty (F(1,24) = 2.71, p = 0.11). There was an interaction between previous difficulty × ROI (F(6,144) = 12.29, p < 0.001), as well as current difficulty × ROI (F(6,144) = 10.13, p < 0.001), but no previous difficulty × current difficulty interaction (F(1,24) = 1.88, p = 0.18). There was no previous difficulty × current difficulty × ROI interaction (F(6,144) = 1.63, p = 0.14). To disentangle the interactions with ROIs, we conducted a previous difficulty × current difficulty ANOVA on each of the MD ROIs. Among the 7 ROIs, the ACC and pdLFC showed a significant main effect for previous difficulty (both Fs(1,24) > 6.17, all ps < 0.03). In ACC, pdLFC, and pMFG there was a main effect of current difficulty (all Fs(1,24) > 7.49, all ps < 0.02). The main effect of previous difficulty was driven by increased activation to the context cue if the previous trial came from the easy set. The main effect of current difficulty was driven by increased activity if the current context cue indicated an upcoming hard math problem. Finally, none of the ROIs showed a main effect of switch type (all Fs(1,24) < 4.21, all ps > 0.05).

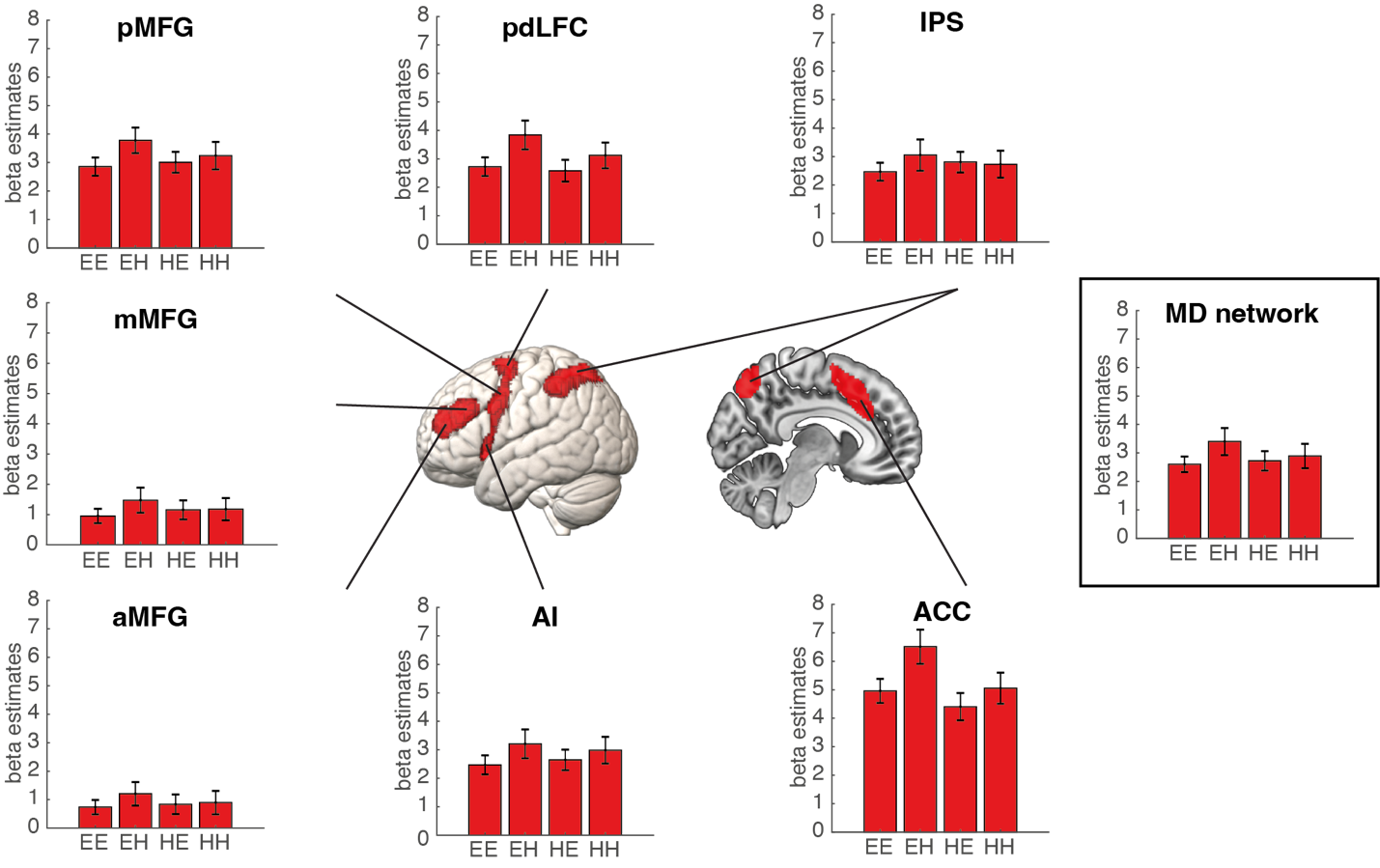

*Figure S1. Activity in the MD network during the context cue based on previous and current trial difficulty. EE: previous easy current easy; EH: previous easy current hard; HE: previous hard current easy; HH: previous hard current hard. Error bars represent standard error.*

#### 1.2. Whole-brain analysis

Results from the whole-brain analysis are presented in Figure S2. During the context cue, MD regions including the ACC and pdLFC, along with regions outside the MD network, including the anterior medial prefrontal cortex, precuneus, posterior cingulate cortex, sensorimotor cortex, visual cortex, and parts of the temporal lobe showed increased activity when the previous context was easy (Figure S2i). Several regions showed a main effect of current difficulty, with MD regions including the ACC and pdLFC as well as the neighboring motor cortex showing an increased response to the doors cueing the hard set. Other areas that showed increased activity to the hard doors included the medial prefrontal cortex, parts of the temporal lobe, parts of the occipital lobe, putamen, basal ganglia, and cerebellum. No region showed increased activity towards the easy doors (Figure S2Ai). We did not find any significant activations for switching to an easy set versus repeating an easy set. However, most of the DMN, along with the motor cortex, posterior insula, temporal lobe, and extrastriatal visual cortex showed increased activity when repeating an easy set versus switching to an easy set (Figure S2Aiii). Switching to a hard set versus repeating a hard set activated large swathes of the DMN, along with some MD regions, including the ACC and pdLFC, as well as the motor cortex, visual cortex, and temporal lobe. No regions showed increased activity when repeating a hard set (Figure S2Aiv).

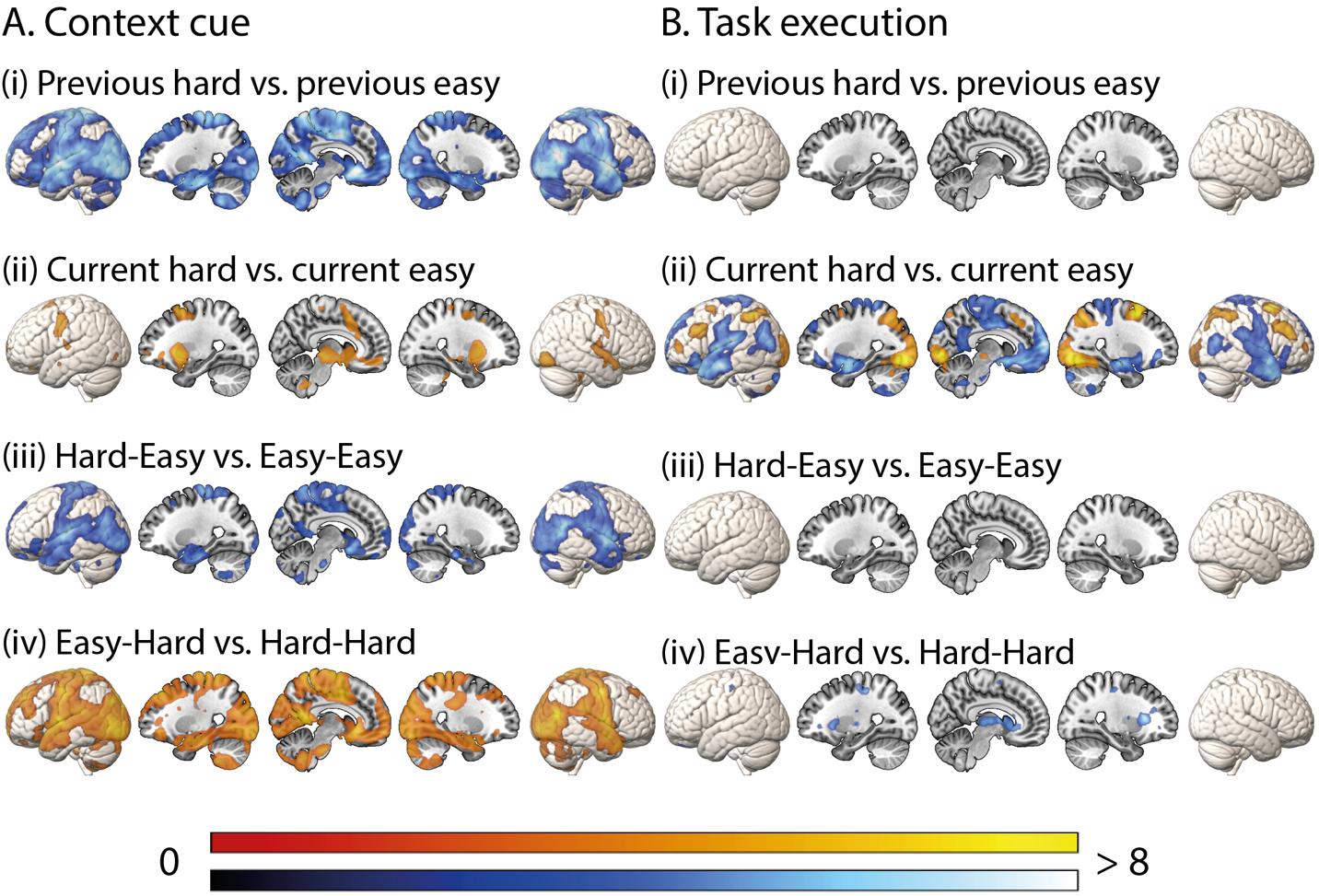

*Figure S2. Whole-brain univariate analysis during the context cue (A) and task execution (B), showing (i) previously hard versus easy set, (ii) current hard versus easy set, (iii) switching to an easy set versus repeating an easy set, and (iv) switching to a hard set versus repeating a hard set. Univariate results during task execution (B) showing hard versus easy problems. Colors indicate t values, with warm and cool scales indicating positive and negative tails, respectively. The activation maps are thresholded at FDR < 0.025 per tail.*

### 2. Corrected and uncorrected statistics per ROI

#### 2.1. Univariate activation across difficulty conditions

The Tables below show descriptive statistics for pairwise t-tests across the 6 experimental conditions (Easy-Low, Easy-Medium, Easy-High, Hard-Low, Hard-Medium, and Hard-High) for each individual MD ROI. Reported p values are FDR corrected across ROIs and conditions. Parathesis contain the uncorrected p values.

aMFG

|  | Easy-Medium | Easy-High | Hard-Low | Hard-Medium | Hard-High |
| --- | --- | --- | --- | --- | --- |
| Easy-Low | t = 1.08  p = 0.41 (0.29) | t = 0.13  p = 0.94 (0.90) | t = 0.25  p = 0.88 (0.81) | t = -0.53  p = 0.73 (0.60) | t = -0.18  p = 0.91 (0.86) |
| Easy-Medium |  | t = -1.87  p = 0.14 (0.07) | t = -1.46  p = 0.25 (0.16) | t = -3.63  p = 0.01 (0.001) | t = -2.49  p = 0.05 (0.02) |
| Easy-High |  |  | t = 0.28  p = 0.88 (0.78) | t = -1.64  p = 0.20 (0,11) | t = -0.64  p = 0.67 (0.52) |
| Hard-Low |  |  |  | t = -1.64  p = 0.20 (0.11) | t = -0.66  p = 0.66 (0.52) |
| Hard-Medium |  |  |  |  | t = 0.68  p = 0.65 (0.50) |

mMFG

|  | Easy-Medium | Easy-High | Hard-Low | Hard-Medium | Hard-High |
| --- | --- | --- | --- | --- | --- |
| Easy-Low | t = -0.73  p = 0.62 (0.47) | t = -2.90  p = 0.02 (0.01) | t = -2.33  p = 0.07 (0.03) | t = -3.17  p = 0.01 (<0.01) | t = -3.30  p = 0.01 (<0.01) |
| Easy-Medium |  | t = -3.84  p < 0.01 (<0.001) | t = -3.73  p = 0.01 (0.001) | t = -7.00  p < 0.001 (<0.001) | t = -5.39  p < 0.001 (<0.001) |
| Easy-High |  |  | t = 0.23  p = 0.88 (0.82) | t = -2.01  p = 0.12 (0.06) | t = -1.75  p = 0.17 (0.09) |
| Hard-Low |  |  |  | t = -2.33  p = 0.07 (0.03) | t = -1.41  p = 0.26 (0.17) |
| Hard-Medium |  |  |  |  | t = 0.22  p = 0.88 (0.83) |

pMFG

|  | Easy-Medium | Easy-High | Hard-Low | Hard-Medium | Hard-High |
| --- | --- | --- | --- | --- | --- |
| Easy-Low | t = -2.55  p < 0.05 (0.02) | t = -4.46  p < 0.01 (< 0.001) | t = -4.03  p < 0.01 (< 0.001) | t = -4.40  p < 0.01 (< 0.001) | t = -4.16  p < 0.01 (< 0.001) |
| Easy-Medium |  | t = -3.53  p < 0.01 (< 0.01) | t = -2.80  p = 0.03 (0.01) | t = -5.29  p < 0.001 (< 0.001) | t = -3.74  p < 0.01 (0.001) |
| Easy-High |  |  | t = 0.72  p = 0.63 (0.48) | t = -1.08  p = 0.41 (0.29) | t = -0.61  p = 0.69 (0.55) |
| Hard-Low |  |  |  | t = -2.02  p = 0.11 (0.05) | t = -1.01  p = 0.45 (0.32) |
| Hard-Medium |  |  |  |  | t = 0.27  p = 0.88 (0.79) |

pdLFC

|  | Easy-Medium | Easy-High | Hard-Low | Hard-Medium | Hard-High |
| --- | --- | --- | --- | --- | --- |
| Easy-Low | t = -1.98  p = 0.12 (0.06) | t = -3.98  p < 0.01 (< 0.001) | t = -2.81  p = 0.03 (0.01) | t = -4.06  p < 0.01 (< 0.001) | t = -3.57  p < 0.01 (< 0.01) |
| Easy-Medium |  | t = -2.89  p = 0.02 (< 0.01) | t = -1.61  p = 0.20 (0.12) | t = -4.10  p < 0.01 (< 0.001) | t = -3.57  p < 0.01 (< 0.01) |
| Easy-High |  |  | t = 1.19  p = 0.35 (0.25) | t = -1.48  p = 0.24 (0.15) | t = -0.76  p = 0.61 (0.45) |
| Hard-Low |  |  |  | t = -3.02  p = 0.02 (< 0.01) | t = -1.46  p = 0.25 (0.16) |
| Hard-Medium |  |  |  |  | t = 0.38  p = 0.85 (0.70) |

IPS

|  | Easy-Medium | Easy-High | Hard-Low | Hard-Medium | Hard-High |
| --- | --- | --- | --- | --- | --- |
| Easy-Low | t = -2.96  p = 0.02 (<0.01) | t = -4.71  p = 0.001 (< 0.001) | t = -4.15  p < 0.01 (< 0.001) | t = -5.71  p < 0.001 (< 0.001) | t = -5.25  p < 0.001 (< 0.001) |
| Easy-Medium |  | t = -3.36  p = 0.01 (< 0.01) | t = -3.24  p = 0.01 (< 0.01) | t = -6.00  p < 0.001 (< 0.001) | t = -4.59  p = 0.001 (< 0.001) |
| Easy-High |  |  | t = 0.02  p = 0.98 (0.98) | t = -2.90  p = 0.02 (< 0.01) | t = -2.14  p = 0.09 (0.04) |
| Hard-Low |  |  |  | t = -3.24  p = 0.01 (< 0.01) | t = -1.88  p = 0.14 (0.07) |
| Hard-Medium |  |  |  |  | t = 0.24  p = 0.88 (0.82) |

AI

|  | Easy-Medium | Easy-High | Hard-Low | Hard-Medium | Hard-High |
| --- | --- | --- | --- | --- | --- |
| Easy-Low | t = -0.11  p = 0.94 (0.91) | t = -1.98  p = 0.12 (0.06) | t = -1.63  p = 0.20 (0.12) | t = -2.49  p = 0.05 (0.02) | t = -2.27  p = 0.08 (0.03) |
| Easy-Medium |  | t = -2.97  p = 0.02 (< 0.01) | t = -2.23  p = 0.08 (0.04) | t = -5.18  p < 0.001 (< 0.001) | t = -4.77  p < 0.001 (< 0.001) |
| Easy-High |  |  | t = -0.11  p = 0.94 (0.92) | t = -1.91  p = 0.13 (0.07) | t = -1.63  p = 0.20 (0.12) |
| Hard-Low |  |  |  | t = -1.35  p = 0.28 (0.19) | t = -0.88  p = 0.53 (0.39) |
| Hard-Medium |  |  |  |  | t = 0.06  p = 0.97 (0.95) |

ACC

|  | Easy-Medium | Easy-High | Hard-Low | Hard-Medium | Hard-High |
| --- | --- | --- | --- | --- | --- |
| Easy-Low | t = -0.54  p = 0.73 (0.59) | t = -1.57  p = 0.21 (0.13) | t = -1.43  p = 0.26 (0.17) | t = -1.35  p = 0.28 (0.19) | t = -1.34  p = 0.28 (0.19) |
| Easy-Medium |  | t = -2.27  p = 0.08 (0.03) | t = -1.64  p = 0.20 (0.11) | t = -2.25  p = 0.08 (0.03) | t = -2.04  p = 0.11 (0.05) |
| Easy-High |  |  | t = 0.28  p = 0.88 (0.78) | t = 0.05  p = 0.97 (0.96) | t = -0.26  p = 0.88 (0.80) |
| Hard-Low |  |  |  | t = -0.24  p = 0.88 (0.81) | t = -0.37  p = 0.85 (0.72) |
| Hard-Medium |  |  |  |  | t = -0.36  p = 0.85 (0.72) |

#### 2.2. Univariate activation when switching difficulty contexts

The Tables below show descriptive statistics for the previous difficulty (easy vs. hard) × current difficulty (easy vs. hard) ANOVA performed on each individual MD ROI during the context cue and task execution, respectively. Reported p values are FDR corrected across ROIs. Parathesis contain the uncorrected p values.

##### 2.2.1. Context Cue

|  | previous | current | previous × current |
| --- | --- | --- | --- |
| aMFG | F(1,24) = 0.30  p = 0.97 (0.59) | F(1,24) = 1.31  p = 0.31 (0.26) | F(1,24) = 0.69  p = 0.42 (0.41) |
| mMFG | F(1,24) = 0.05  p = 0.97 (0.82) | F(1,24) = 1.78  p = 0.27 (0.19) | F(1,24) = 1.31  p = 0.37 (0.26) |
| pmFG | F(1,24) = 1..35  p = 0.60 (0.26) | F(1,24) = 7.50  p = 0.03 (0.01) | F(1,24) = 4.21  p = 0.29 (0.05) |
| pdLFC | F(1,24) = 6.17  p = 0.07 (0.02) | F(1,24) = 11.99  p < 0.01 (< 0.01) | F(1,24) = 1.97  p = 0.30 (0.17) |
| IPS | F(1,24) < 0.01  p = 0.97 (0.97) | F(1,24) = 0.90  p = 0.35 (0.35) | F(1,24) = 3.03  p = 0.29 (0.09) |
| AI | F(1,24) = 0.03  p = 0.97 (0.86) | F(1,24) = 3.51  p = 0.13 (0.07) | F(1,24) = 0.67  p = 0.42 (0.42) |
| ACC | F(1,24) = 26.58  p < 0.001 (< 0.001) | F(1,24) = 16.09  p < 0.01 (< 0.001) | F(1,24) = 2.52  p = 0.29 (0.13) |

##### 2.2.2. Task Execution

|  | previous | current | previous × current |
| --- | --- | --- | --- |
| aMFG | F(1,24) = 0.45  p = 0.71 (0.51) | F(1,24) = 3.52  p = 0.09 (0.07) | F(1,24) = 0.24  p = 0.74 (0.63) |
| mMFG | F(1,24) = 0.07  p = 0.93 (0.80) | F(1,24) = 22.97  p < 0.001 (< 0.001) | F(1,24) = 0.26  p = 0.74 (0.62) |
| pmFG | F(1,24) = 1.40  p = 0.58 (0.25) | F(1,24) = 14.33  p < 0.01 (< 0.001) | F(1,24) = 0.25  p = 0.74 (0.62) |
| pdLFC | F(1,24) = 2.46  p = 0.57 (0.13) | F(1,24) = 6.80  p = 0.02 (0.02) | F(1,24) = 0.26  p = 0.74 (0.62) |
| IPS | F(1,24) < 0.01  p = 0.96 (0.96) | F(1,24) = 27.63  p < 0.001 (< 0.001) | F(1,24) = 0.58  p = 0.74 (0.45) |
| AI | F(1,24) = 2.06  p = 0.57 (0.16) | F(1,24) = 17.59  p < 0.001 (< 0.001) | F(1,24) < 0.01  p = 0.96 (0.96) |
| ACC | F(1,24) = 0.77  p = 0.68 (0.39) | F(1,24) = 0.64  p = 0.43 (0.43) | F(1,24) = 0.29  p = 0.74 (0.59) |

#### 2.3. RSA regression

The Table below shows the t statistics for beta coefficients (context-independent, context-dependent, and pixelwise dissimilarity) tested against 0 (1-tailed) on each individual MD ROI. Reported p values are FDR corrected across ROIs. Parathesis contain the uncorrected p values.

|  | context-independent | context-dependent | pixelwise dissimilarity |
| --- | --- | --- | --- |
| aMFG | t = 7.75  p < 0.001 (< 0.001) | t = 1.73  p = 0.22 (0.16) | t = 1.33  p = 0.22 (0.16) |
| mMFG | t = 5.32  p < 0.001 (< 0.001) | t = 2.09  p = 0.16 (<0.05) | t = 0.02  p = 0.21 (0.10) |
| pmFG | t = 10.18  p < 0.001 (< 0.001) | t = 0.39  p = 0.17 (0.10) | t = -0.02  p = 0.21 (0.09) |
| pdLFC | t = 9.68  p < 0.001 (< 0.001) | t = -0.81  p = 0.79 (0.79) | t = 1.22  p = 0.21 (0.12) |
| IPS | t = 24.74  p < 0.001 (< 0.001) | t = 1.55  p = 0.16 (0.07) | t = 1.56  p = 0.21 (0.07) |
| AI | t = 4.91  p < 0.001 (< 0.001) | t = 1.02  p = 0.16 (0.02) | t = 1.02  p = 0.51 (0.49) |
| ACC | t = 12.28  p < 0.001 (< 0.001) | t = 1.33  p = 0.41 (0.35) | t = 1.40  p = 0.51 (0.51) |

#### 2.4. LDC distances for matched conditions

The Table below shows the t statistics for LDC distances between the high difficulty level in the easy set and the low difficulty level in the hard set tested against 0 (1-tailed) on each individual MD ROI. Reported p values are FDR corrected across ROIs. Parathesis contain the uncorrected p values.

|  | Descriptive statistics |
| --- | --- |
| aMFG | t = -0.54  p = 0.98 (0.70) |
| mMFG | t = -1.37  p = 0.98 (0.91) |
| pmFG | t = -1.52  p = 0.98 (0.93) |
| pdLFC | t = -0.24  p = 0.98 (0.59) |
| IPS | t = -2.09  p = 0.98 (0.98) |
| AI | t = 1.05  p = 0.98 (0.15) |
| ACC | t = 0.50  p = 0.98 (0.31) |

### 3. Pixelwise dissimilarity

For each participant, we constructed an additional model RDM estimating the pixel-level correlation distance between the mean math stimuli of the different conditions as they were presented on screen. The average pixel RDM across participants is illustrated in Figure S3. We note that the pixel-level correlation distance between difficulty levels do not linearly correspond to objective difficulty levels of the task, and this is due to the center alignment of the stimuli, such that distance is influenced by both the symmetry of the stimuli as well as number of black pixels on the screen.

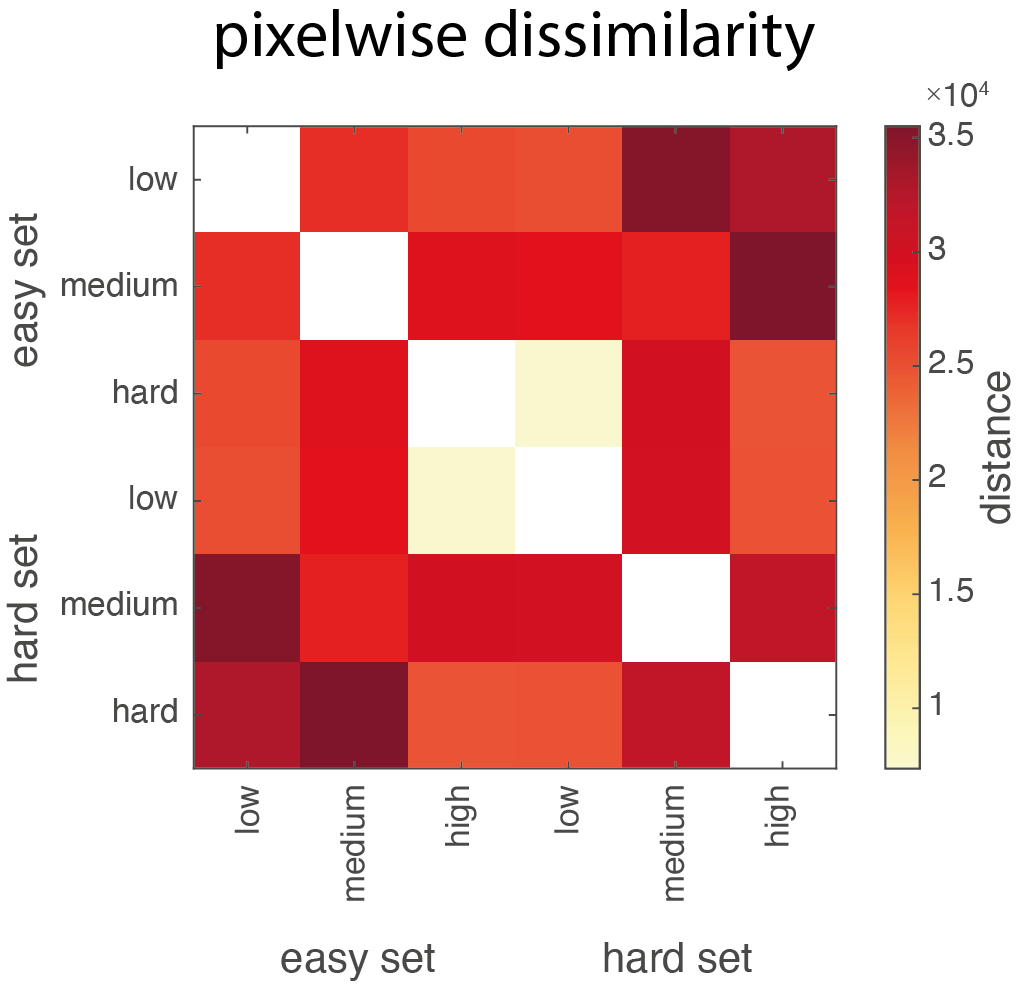

*Figure S3. RDM of average pixelwise dissimilarity between the six conditions.*

### 4. Empirical RDMs of MD regions

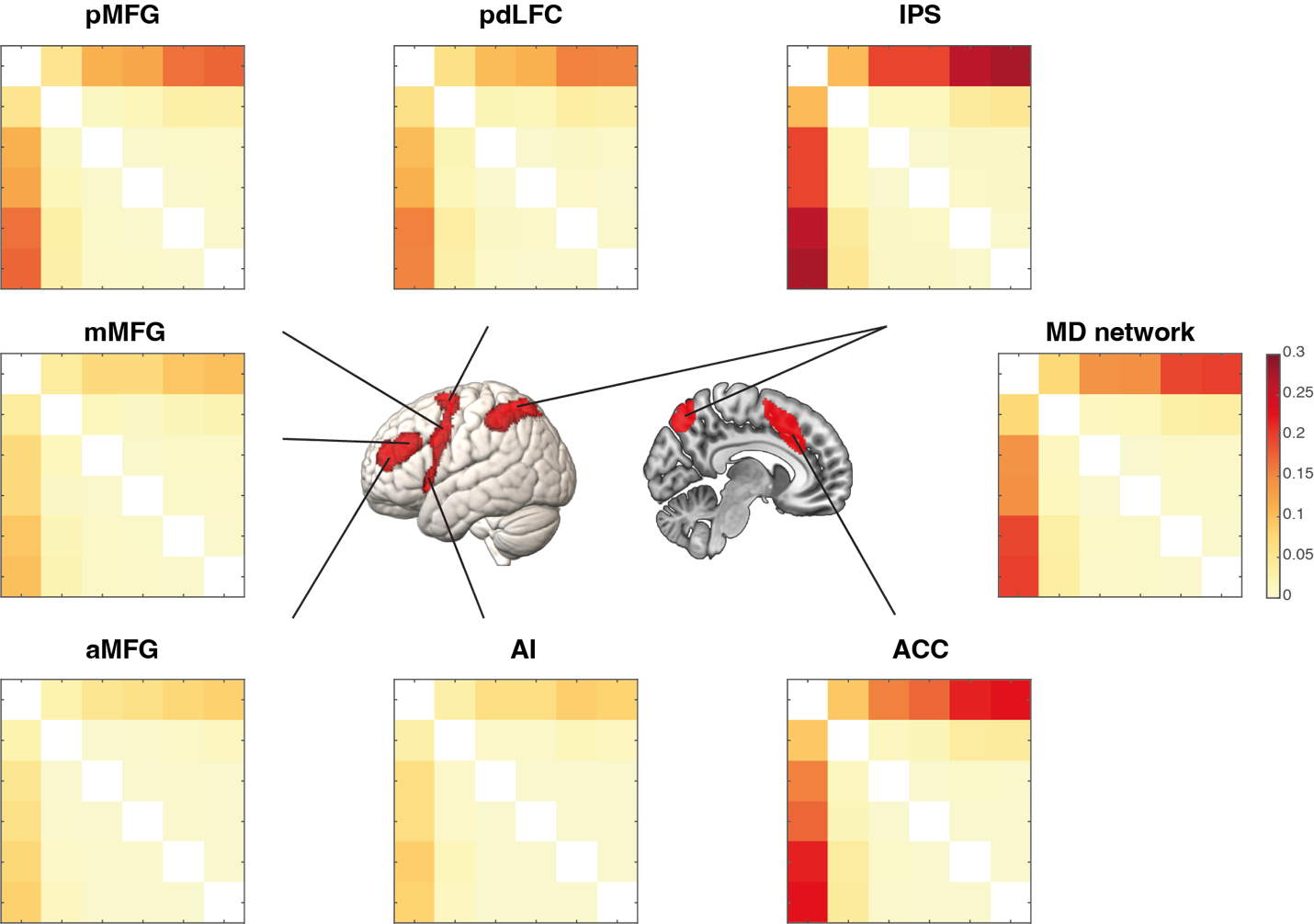

*Figure S4. Empirical RDMs for each MD ROI. The colors indicate LDC distances normalized by the number of voxels within each ROI, with warmer colors indicating greater distance.*

### 5. Analysis of the DMN

#### 5.1. ROIs

We used DMN ROIs from Wen et al. (2020), which were constructed using the 17 network parcellation from Yeo et al. (2011) and divided into ROIs using coordinates described in Andrews-Hanna et al. (2010). Details of these ROIs can be found in Wen et al. (2020). Bilateral ROIs were combined as our previous study suggest they show similar activity patterns and task representations (Wen et al. 2020). This resulted in a total of 11 ROIs, including the dorsal medial prefrontal cortex (dMPFC), temporal parietal junction (TPJ), lateral temporal cortex (LTC), temporal pole (TempP), ventral medial prefrontal cortex (vMPFC), posterior inferior parietal lobule (pIPL), retrosplenial cortex (Rsp), parahippocampal cortex (PHC), hippocampal formation (HF+), anterior medial prefrontal cortex (aMPFC), and posterior cingulate cortex (PCC).

#### 5.2. Univariate activation across difficulty conditions

Average beta estimates of each difficulty level from bilateral DMN regions, as well as a combined DMN network ROI, are shown in Figure S5. We first ran a context (easy vs. hard) × difficulty level (low, medium, and high) repeated measures ANOVA to examine activity in the DMN network. Results showed a significant main effect of context (F(1,24) = 23.48, p < 0.001) and a significant main effect of difficulty level (F(2,48) = 9.59, p < 0.001). There was also a context × difficulty level interaction (F(2,48) = 13.29, p < 0.001). Pairwise t-tests across the six difficulty conditions revealed showed that the low difficulty condition in the easy set was significantly different from all other conditions (all ts > 3.82, all ps < 0.001). There were no significant differences between any of the other difficulty conditions (all |t|s < 1.54, all ps > 0.13).

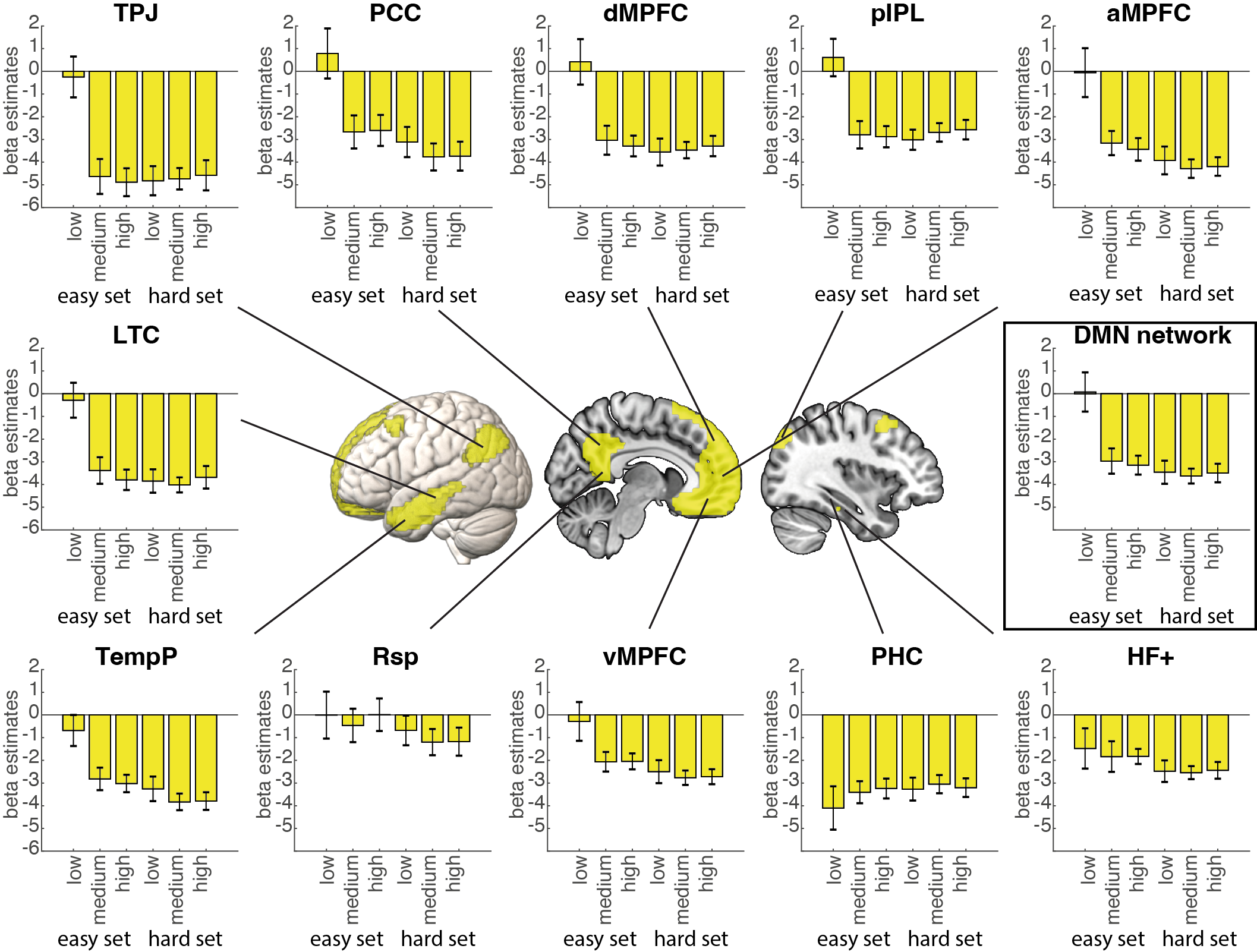

*Figure S5. ROI results of DMN regions (left) as well as the entire DMN network (right). Graphs show the beta values for each math condition. Error bars represent standard error.*

To compare across difficulty levels and ROIs, we conducted a 3-way ANOVA with factors context (easy vs. hard), difficulty level (low, medium, and high), and ROI (11 DMN ROIs). This analysis showed a significant main effect of context (F(1,24) = 16.35, p < 0.001), a significant main effect of difficulty level (F(2,48) = 7.26, p < 0.01), and a significant main effect of ROI (F(10,240) = 9.25, p < 0.001). The main effect of context was driven by decreased DMN activity in the hard compared to easy context, and the main effect of difficulty was driven by decreasing DMN activity with higher difficulty levels. There was a context × ROI interaction (F(10,240) = 7.05, p < 0.001), difficulty level × ROI interaction (F(20,480) = 7.51, p < 0.001), as well as context × difficulty level interaction (F(2,48) = 7.47, p < 0.01). Finally, there was a context × difficulty level × ROI interaction (F(20,480) = 9.62, p < 0.001). Pairwise t-tests across the six difficulty conditions revealed that in all of the DMN ROIs, except for the Rsp, PHC, and HF+, there was a decrease in activation and a later as plateau as difficulty increased. The sharpest decrease was seen between the low and medium difficulty levels in the easy condition. There were no differences in activation among the low, medium, and high difficulty levels in the hard condition in any of the ROIs (max t = 1.93, p = 0.19).

We also examined the two conditions matched in absolute difficulty, the high difficulty level in the easy context and the low difficulty level in the hard context. We found no difference in the DMN network ROI in response to these two conditions (t = 0.84, p = 0.40). Furthermore, none of the individual ROIs showed any significant differences between the matched difficulty conditions after FDR correction (all |t|s < 2.11, all ps > 0.50). Without multiple comparison correction, HF+ showed lower activation in the hard set high difficulty level than the easy set low difficulty level condition (t = 2.10, p < 0.05).

#### 5.3. Univariate activation when switching difficulty contexts

For the context cue and task execution, we separately performed a previous difficulty (easy vs. hard) × current difficulty (easy vs. hard) ANOVA on the combined DMN ROI. To further examine differences between ROIs, we also conducted a previous difficulty (easy vs. hard) × current (easy vs. hard) × ROI (11 DMN ROIs) ANOVA.

##### 5.3.1. Context cue

For activity during the context cue presentation (Figure S6), the DMN network ROI showed a main effect of previous difficulty (F(1,24) = 20.47, p < 0.001), but no main effect of current difficulty (F(1,24) = 1.65, p = 0.21), and no previous difficulty × current difficulty interaction (F(1,24) = 0.16, p = 0.69). The main effect of previous difficulty was driven by higher DMN activity when the previous trial was from the easy set compared to hard set. Adding the 11 ROIs as an additional factor of interest, the ANOVA results showed a main effect of previous difficulty (F(1,24) = 31.90, p < 0.001), a main effect of ROI (F(10,240) = 9.44, p < 0.001), but no main effect of current difficulty (F(1,24) = 2.43, p = 0.13). The main effect of previous difficulty again was driven by higher DMN activity when the previous trial was from the easy set compared to hard set. There was an interaction between previous difficulty × ROI (F(10,240) = 4.77, p < 0.001), as well as current difficulty × ROI (F(10,240) = 2.84, p = 0.02), but no previous difficulty × current difficulty interaction (F(1,24) = 0.23, p = 0.64). Finally, there was a previous difficulty × current difficulty × ROI interaction (F(10,240) = 2.45, p = 0.03).

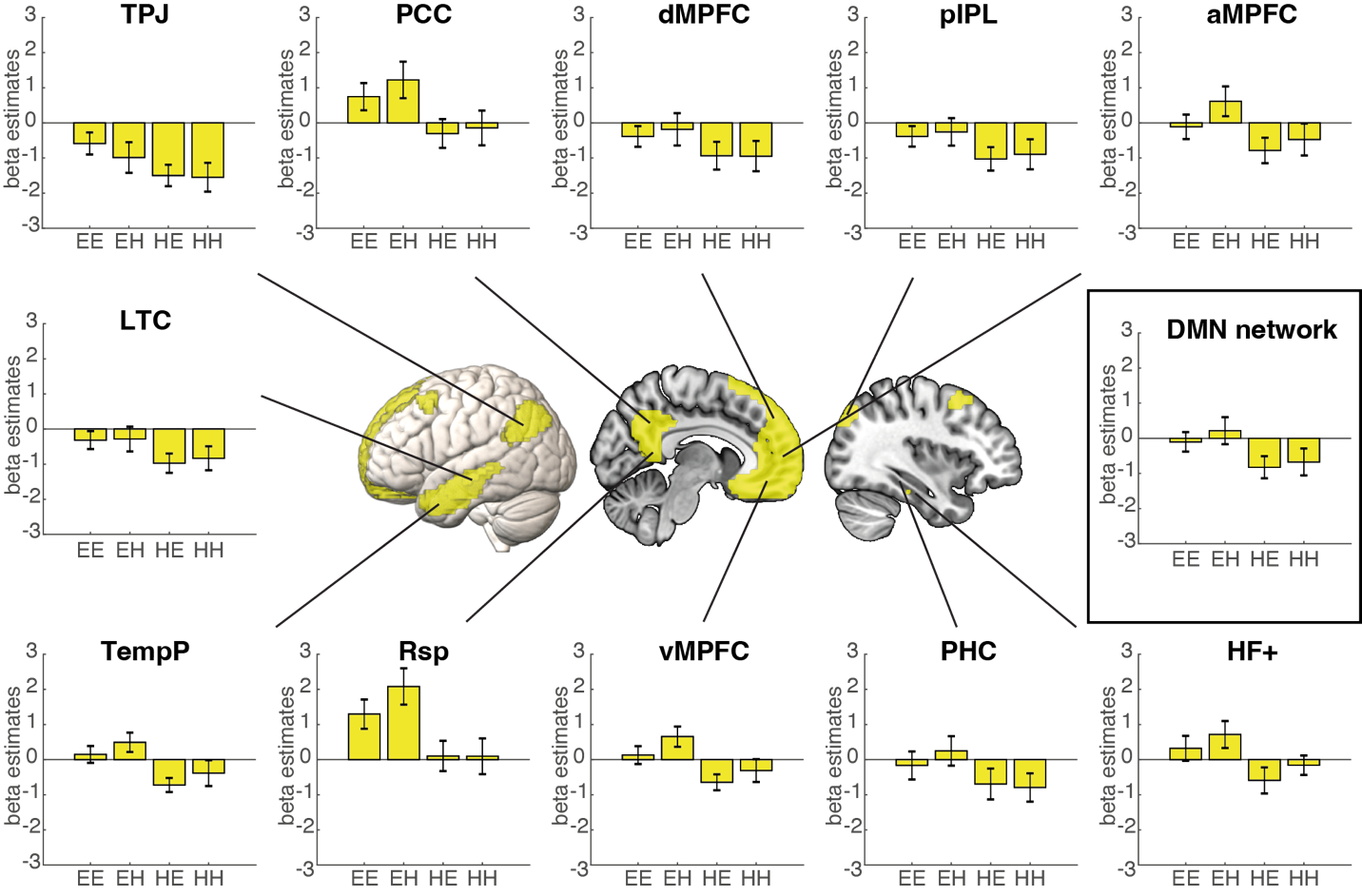

*Figure S7. Activity in the DMN during the context cue based on previous and current trial difficulty. EE: previous easy current easy; EH: previous easy current hard; HE: previous hard current easy; HH: previous hard current hard. Error bars represent standard error.*

##### 5.3.2. Task execution

For activity during task execution (Figure S7), the DMN network ROI showed a main effect of current difficulty (F(1,24) = 16.28, p < 0.001), but no main effect of previous difficulty (F(1,24) = 0.19, p = 0.67), and no previous difficulty × current difficulty interaction (F(1,24) = 0.01, p = 0.94). The main effect of current difficulty was driven by higher DMN activity when the current trial was from the easy set compared to hard set. Adding the 11 ROIs as an additional factor of interest, the ANOVA results showed a main effect of current difficulty (F(1,24) = 12.40, p < 0.01), a main effect of ROI (F(10,240) = 12.26, p < 0.001), but no main effect of previous difficulty (F(1,24) = 0.46, p = 0.51). The main effect of current difficulty the result of higher DMN activity when the current trial was from the easy set compared to hard set. There was an interaction between current difficulty × ROI (F(10,240) = 6.04, p < 0.001), but no previous difficulty × ROI (F(10,240) = 1.14, p = 0.34), previous difficulty × current difficulty interaction (F(1,24) = 0.07, p = 0.79), and no previous difficulty × current difficulty × ROI interaction (F(10,240) = 0.74, p = 0.57).

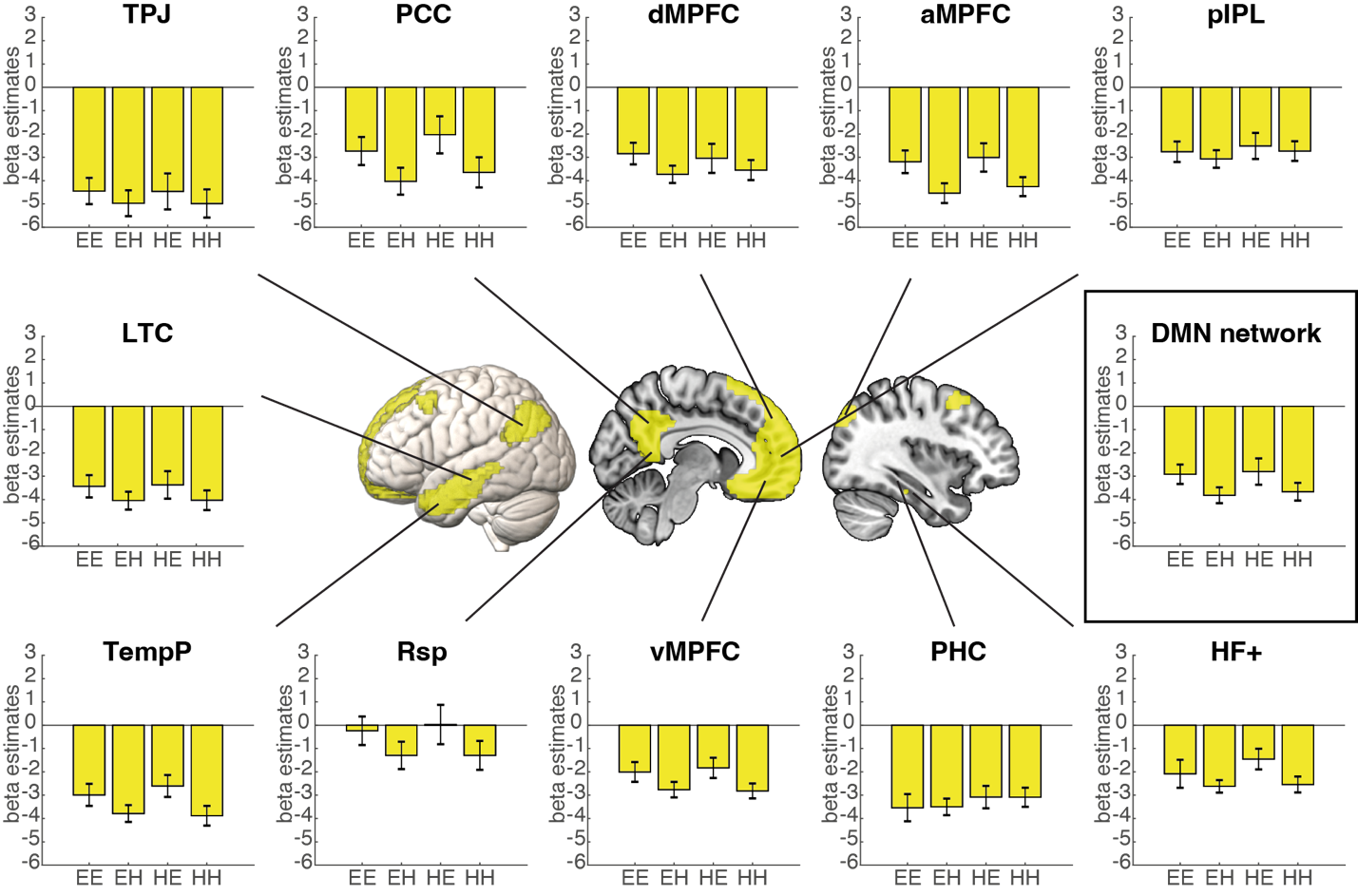

*Figure S8. Activity in the DMN during the task execution based on previous and current trial difficulty. EE: previous easy current easy; EH: previous easy current hard; HE: previous hard current easy; HH: previous hard current hard. Error bars represent standard error.*

In summary, these results suggest that the DMN may be sensitive to difficulty context during cue processing; however, during task execution, we were only able to identify DMN sensitivity towards the current and not previous difficulty context. We also note that it while the effect of previous difficulty context during the cue phase may indicate that the DMN is sensitive to temporal context, this may also reflect carryover effects from the previous trial, for example, participants may be still thinking about the previous math problem during the cue phase of the current trial. Future studies may be needed to disentangle these two possibilities.

#### 5.4. RSA results

Results are shown in Figure S9 and empirical RDMs are shown in Figure S10. In DMN ROIs, the beta coefficients of the context-independent model RDM were significantly greater than zero in all ROIs (all ts > 2.85, all ps < 0.01; FDR corrected) except for the TempP (t = 1.39, p = 0.09; FDR corrected), indicating a relationship with the brain RDMs. The context-independent model RDM provided a significantly better fit than the context-dependent model RDM in all DMN ROIs (all ts > 1.95, all ps < 0.04; FDR corrected) except for the TempP (t = 1.49, p = 0.07; FDR corrected). The coefficients of the context-dependent model RDM were significantly above zero in many of the DMN ROIs, including the dMPFC, LTC, TPJ, and Rsp (all ts > 2.30, all ps < 0.04; FDR corrected). We also note that the coefficients of the context-dependent model RDM in the vMPFC was also significant before multiple comparison correction (t = 2.09, p = 0.02 uncorrected). Finally, the beta coefficients for the pixel RDM were not significantly above zero in any of the ROIs (all |t|s < 1.76, all ps > 0.33; FDR corrected). For the combined DMN network ROI, the beta coefficients of the context-independent model RDM was significantly above zero (t = 17.23, p < 0.001) and had a significantly better fit than the context-dependent model RDM (t = 10.08, p < 0.001), which was significantly above zero (t = 2.03, p = 0.03). The beta coefficients for the pixel RDM were not significantly above zero (t = 1.19, p = 0.12). In summary, our results suggest that the DMN shows context-independent coding for task difficulty, however, some DMN regions may additionally show context-dependent coding task as well.

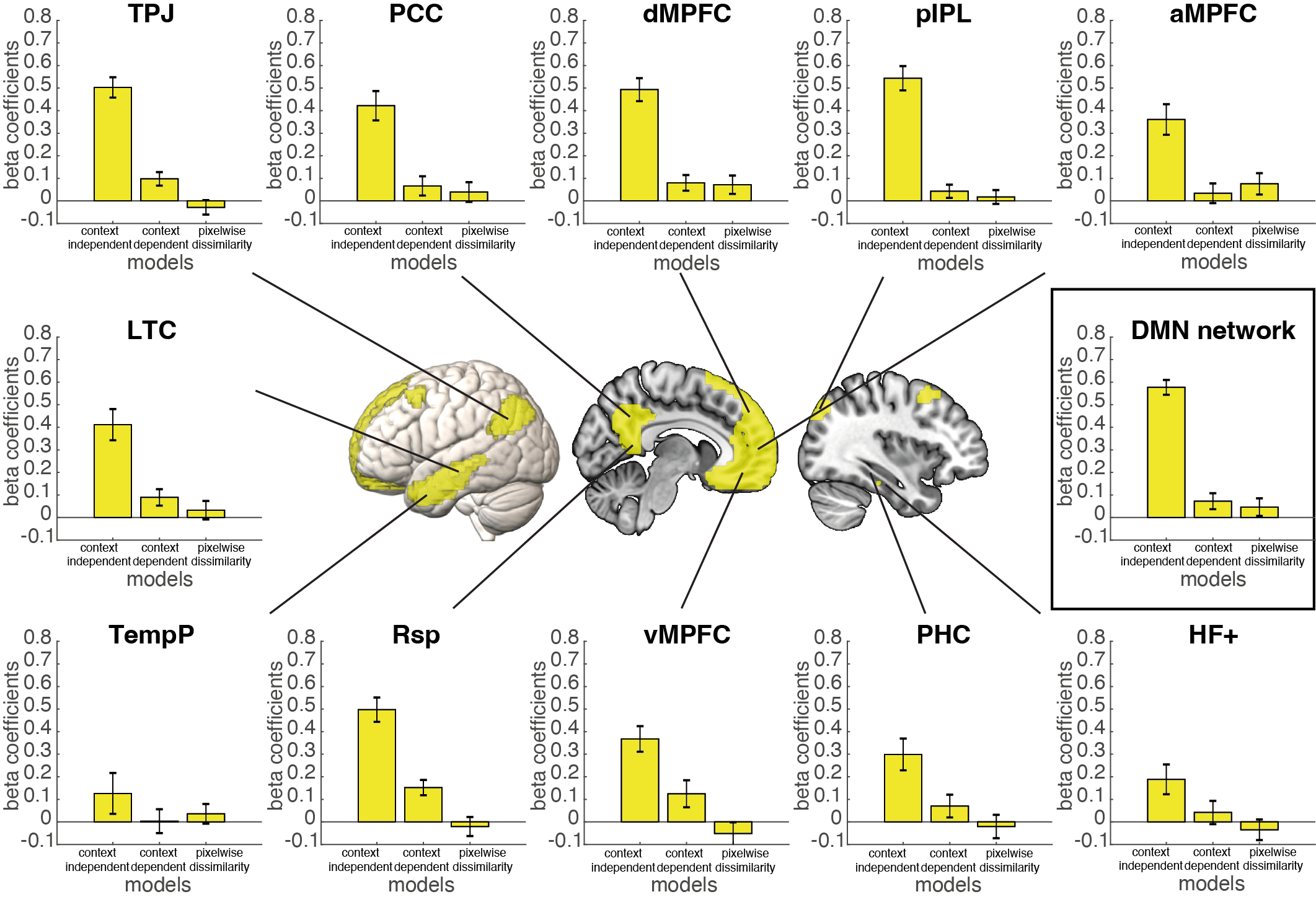

*Figure S9. Relatedness of the model RDMs to the brain RDM for the DMN ROIs (left) and the entire DMN network (right). Beta coefficients were calculated by fitting the three model RDMs (context-independent, context-dependent, and pixelwise dissimilarity) to the brain RDM for each ROI after rank transformation. Error bars represent standard error.*

#### 5.5. Empirical RDMs of DMN regions

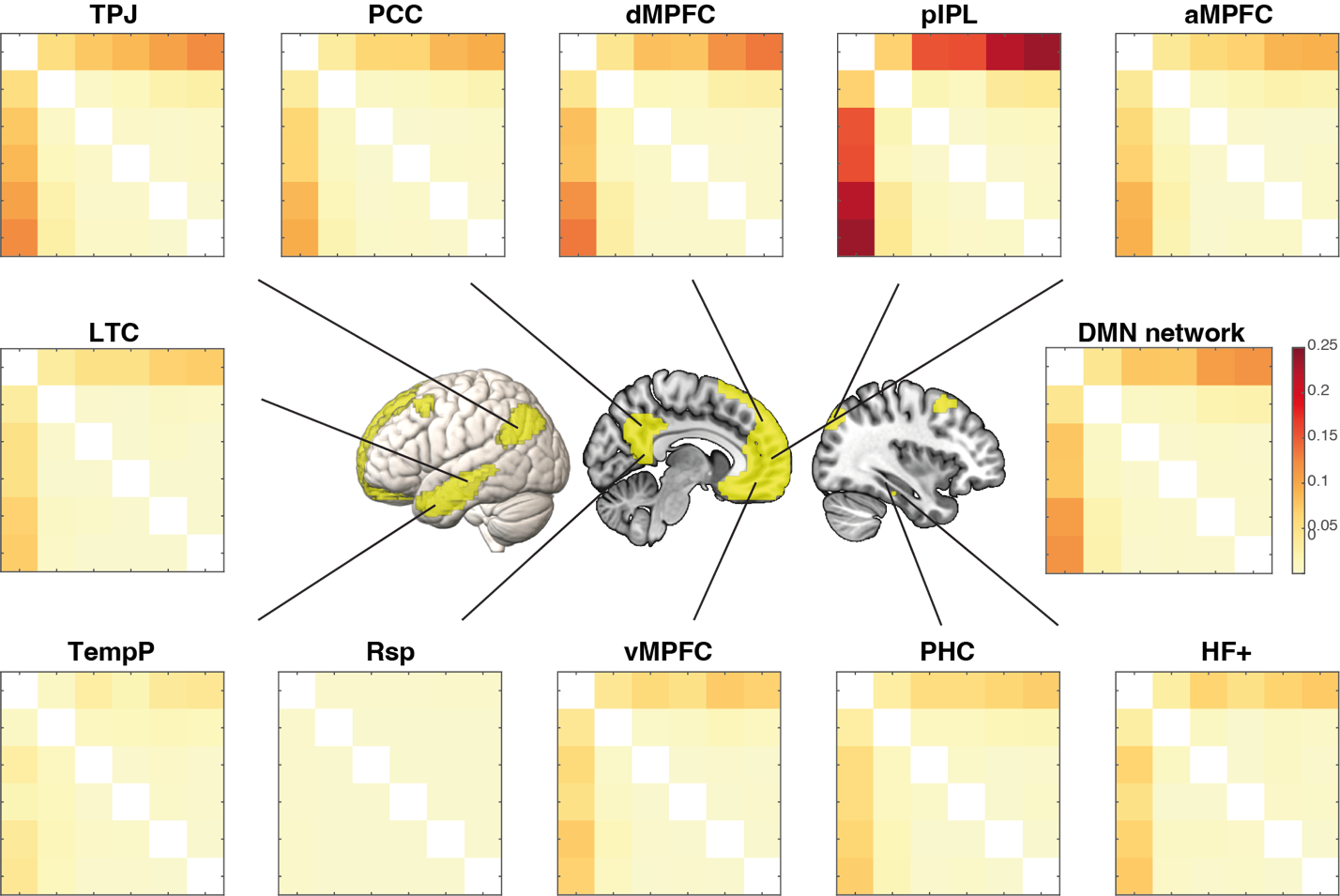

*Figure S10. Empirical RDMs for each DMN ROI. The colors indicate LDC distances normalized by the number of voxels within each ROI, with warmer colors indicating greater distance.*
